# CLYBL averts methylmalonyl-CoA mutase inhibition and loss of vitamin B12 by repairing malyl-CoA

**DOI:** 10.1101/2024.09.25.614871

**Authors:** Corey M. Griffith, Jean-François Conrotte, Parisa Paydar, Xinqiang Xie, Ursula Heins-Marroquin, Floriane Gavotto, Christian Jäger, Kenneth W. Ellens, Carole L. Linster

## Abstract

Citrate lyase beta-like protein (CLYBL) is a ubiquitously expressed mammalian enzyme known for its role in the degradation of itaconate, a bactericidal immunometabolite produced in activated macrophages. The association of *CLYBL* loss-of-function with reduced circulating vitamin B12 levels was proposed to result from inhibition of the B12-dependent enzyme methylmalonyl-CoA mutase (MCM) by itaconyl-CoA. The discrepancy between the highly inducible and locally confined production of itaconate and the broad expression profile of *CLYBL* across tissues, suggested a role for this enzyme beyond itaconate catabolism. We discovered that CLYBL additionally functions as a metabolite repair enzyme for malyl-CoA, a side-product of promiscuous TCA cycle enzymes. We found that *CLYBL* knockout cells, accumulating malyl-CoA but not itaconyl-CoA, show decreased levels of adenosylcobalamin and that malyl-CoA is a more potent inhibitor of MCM than itaconyl-CoA. Our work thus suggests that malyl-CoA plays a role in the B12 deficiency observed in individuals with *CLYBL* loss-of-function.

**Graphical Abstract:** 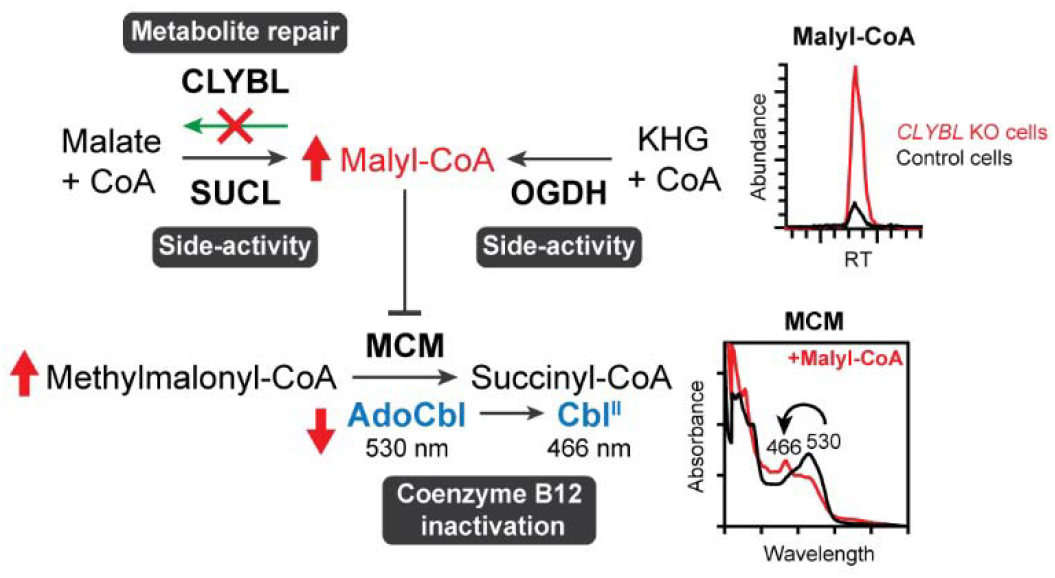

## Introduction

The ubiquitously expressed mitochondrial enzyme citrate lyase beta-like protein (CLYBL) has garnered attention for its role in the catabolism of the antiseptic and anti-inflammatory metabolite, itaconate^1,2^. The latter is formed specifically in activated macrophages by the inducible enzyme IRG1 via decarboxylation of the TCA cycle intermediate *cis*-aconitate, as a defensive mechanism to inhibit a pathogen’s isocitrate lyase^3^. In the host cell, itaconate is successively converted to itaconyl-CoA and citramalyl-CoA, then cleaved into acetyl-CoA and pyruvate by CLYBL^1,2^. Some pathogenic bacteria catabolize itaconate via an analogous pathway involving the bacterial homolog of CLYBL, CitE^1,4^. CitE is a subunit of the heterotrimeric bacterial ATP-independent citrate lyase complex, where it cleaves citryl-CoA into acetyl-CoA and oxaloacetate^5^. In humans, a loss-of-function variant of CLYBL is found with an allele frequency of 2.7% (Genome Aggregation Database^6^) and strongly associated with decreased levels of circulating vitamin B12^7–11^. The corresponding single nucleotide polymorphism (rs41281112) converts Arg259 into a premature stop codon (native CLYBL is 340 amino acids in length), leading to loss of CLYBL protein^2,11^. B12 is an essential nutrient that is processed into methylcobalamin, for methionine synthase to catalyze the conversion of homocysteine to methionine, and adenosylcobalamin, for methylmalonyl-CoA mutase (MCM) to isomerize methylmalonyl-CoA to succinyl-CoA.

Consolidating *in vitro* studies with recombinant human CLYBL^1^, murine brown adipocytes deficient in *CLYBL* were shown to accumulate citramalyl-CoA^2^, confirming that the enzyme acts as a citramalyl-CoA lyase in living cells. The *CLYBL* KO cells also showed increased levels of propionate metabolites, while adenosylcobalamin levels were decreased, indicating that MCM was inhibited and further supporting the connection between CLYBL and B12. Crystallography and spectroscopy studies with human and *Mycobacterium tuberculosis* MCM showed that itaconyl-CoA reacts with the 5’-deoxyadenosyl moiety of the B12 coenzyme to form a biradical adduct that cannot be repaired by the B12 salvage pathway and leads to enzyme inactivation^12^.

Itaconate is not an intermediate of primary metabolism and its formation by the inducible IRG1 enzyme is restricted to activated immune cells^13^. This contrasts with the ubiquitous expression of CLYBL in all tissues^11^, suggesting that the latter not only serves to degrade a metabolite derived from itaconate. Similarly, although itaconyl-CoA clearly intoxicates MCM, the highly inducible and locally confined production of itaconate indicates that it may not be the only cause of reduced circulating B12 levels in *CLYBL* deficient individuals. We therefore hypothesized that CLYBL acts on an additional, more ubiquitously formed acyl-CoA ester through a potential metabolite repair role^14,15^. Toxic or useless metabolites are continuously formed by enzymatic side-activities or spontaneous reactions in primary metabolism, and a devoted set of repair enzymes function to prevent accumulation of these non-canonical metabolites by reverting them to useful or benign products^14,15^. Deficiencies in metabolite repair can cause devastating diseases, including 2-hydroxyglutaric aciduria^16,17^ and the progressive early-onset encephalopathies PEBEL1 and PEBEL2^18,19^. The search for an obscure, ubiquitously formed acyl-CoA ester led us to a substrate of another bacterial enzyme presenting homology with CLYBL/CitE, namely MclA (malyl-CoA lyase)^5^. Malyl-CoA lyases convert malyl-CoA into acetyl-CoA and glyoxylate^20^, operating in the bacterial CO_2_ fixation^21^ and ethylmalonyl-CoA pathways^22^, which are not present in mammals. CLYBL was reported, although not studied further, to have malyl-CoA thioesterase activity *in vitro*^2^, hydrolyzing malyl-CoA into malate and CoA. Intriguingly, *in vitro* studies showed that bacterial succinate-CoA ligase (SUCL) can produce malyl-CoA from malate and CoA^23^. Additionally, the mammalian α-ketoglutarate dehydrogenase complex (henceforth designated KGDH in this article) can convert 2-keto-4-hydroxyglutarate (KHG) and CoA into malyl-CoA *in vitro*^24^. It is thus conceivable that malyl-CoA is formed as a metabolic side product in mammals and that a metabolite repair enzyme is needed to eliminate it.

Herein, we employed enzymatic assays, mammalian cell culture, genome editing, and metabolomics to identify a new, non-canonical mammalian metabolite, malyl-CoA, that is formed by side-activities of TCA cycle enzymes and that is ‘repaired’ by CLYBL to avert inhibition of the B12-dependent enzyme MCM. Importantly, this work uncovered another main physiological function of CLYBL and identifies malyl-CoA as a plausible molecular culprit for reduced circulating vitamin B12 levels in individuals with *CLYBL* deficiency.

## Results

### CLYBL is more efficient as a malyl-CoA thioesterase than a citramalyl-CoA lyase

Malyl-CoA, which is not known as an intermediate of any mammalian metabolic pathway but was reported to be produced by side-activities of TCA cycle enzymes^23,24^, appeared as a good candidate substrate for a putative metabolite repair activity of CLYBL. Enzymatic activity assays of recombinant human CLYBL (Supplementary Fig. 1) revealed a K_*M*_ value of 0.022 mM for its citramalyl-CoA lyase activity (Fig. 1a,b, reaction 1) that is in good agreement with a previously reported value (0.024 mM)^2^. We found an ~10-fold lower turnover number for this activity (*k*_cat_ = 1.6 s^−1^) using our continuous assay compared to the one reported previously (*k*_cat_ = 14.1 s^−1^)^2^ based on an end-point assay (Fig. 1b). We could not measure any malyl-CoA lyase activity with CLYBL using an HPLC-based end-point assay for detection of the expected acetyl-CoA product (Fig 1a, reaction 1). We found, however, that CLYBL actively hydrolyzed malyl-CoA into malate and free CoASH. In fact, CLYBL displayed a 12-fold greater catalytic efficiency as a malyl-CoA thioesterase (855×10^3^ M^−1^.s^−1^, Fig. 1a,b, reaction 2) than as a citramalyl-CoA lyase (72.7×10^3^ M^−1^.s^−1^), strongly supporting that malyl-CoA is a physiological substrate of the human enzyme. Importantly, CLYBL was highly specific as a thioesterase for malyl-CoA compared with other physiological short-chain (C2-C5) acyl-CoA esters (Fig. 1c).

**Fig. 1.**
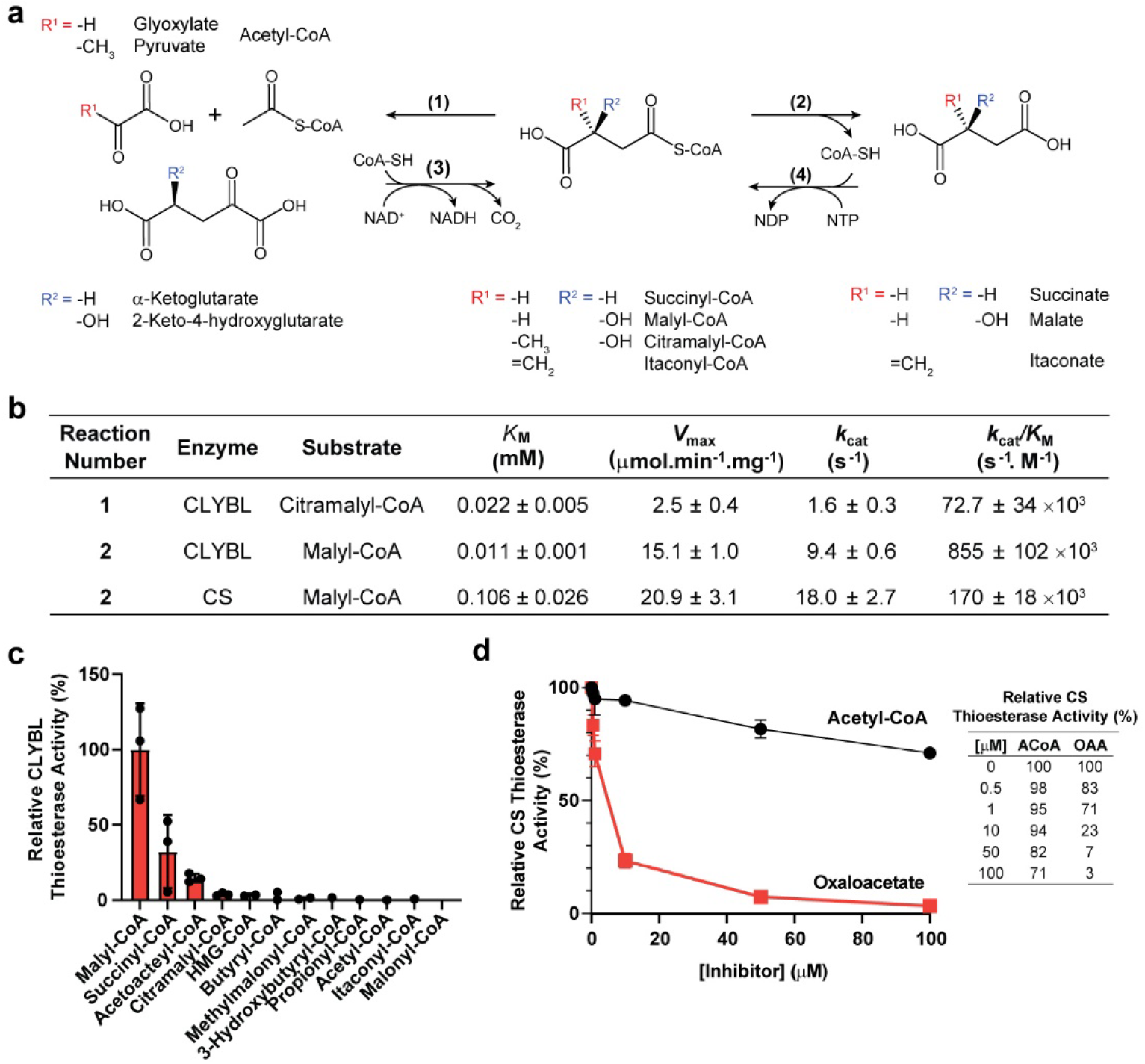
Identification of CLYBL as a malyl-CoA thioesterase. (**a**) Reaction schemes for malyl-CoA/citramalyl-CoA degradation via lyase activity (reaction 1), malyl-CoA degradation via thioesterase activity (reaction 2), and malyl-CoA synthesis starting from 2-keto-4-hydroxyglutarate (KHG; reaction 3) or malate (reaction 4); (**b**) kinetic parameters determined for the citramalyl-CoA lyase activity of CLYBL and the malyl-CoA thioesterase activities of CLYBL and CS, respectively. Malyl-CoA thioesterase activities were measured spectrophotometrically using Ellman’s reagent. Malyl-CoA concentrations between 0-90 μM or 0-700 μM were assayed with CLYBL (0.1 ng/μL) or CS (3 ng/μL), respectively. Citramalyl-CoA lyase activity was measured spectrophotometrically by coupling to lactate dehydrogenase (LDH). Citramalyl-CoA concentration varied between 0-300 μM and CLYBL was added at 2.94 ng/μL; (**c**) substrate specificity of the thioesterase activity of CLYBL (0.1 ng/μL) assayed with the indicated short-chain acyl-CoA esters at a concentration of 100 μM; (**d**) effect of acetyl-CoA (ACoA) or oxaloacetate (OAA) on the malyl-CoA thioesterase activity of CS in the presence of 3 ng/μL CS and 100 μM malyl-CoA. Kinetic parameters were estimated in GraphPad Prism (9.5.0) using the non-linear Michaelis-Menten fit model. The values shown represent means ± SD (n=3). CLYBL, citrate lyase beta-like protein: CS, citrate synthase; HMG-CoA, 3-hydroxy-3-methylglutaryl-CoA.

A malyl-CoA thioesterase activity had previously been reported for porcine heart citrate synthase (CS)^24^ that we compared with the one found here for CLYBL. Using commercial porcine heart CS, we obtained *K*_m_ and *V*_max_ values (0.106 mM and 20.9 μmol.min^−1^.mg^−1^ measured at 37°C; Fig. 1b) that were very close to the reported ones (0.111 mM and 10 μmol.min^−1^.mg^−1^ measured at 30 °C)^24^, corresponding to a catalytic efficiency for malyl-CoA hydrolysis (170×10^3^ s^−1^.M^−1^) that is 5-fold lower compared to the one of CLYBL (Fig. 1b). This indicated that CLYBL is the main enzyme involved in malyl-CoA degradation, a notion further supported by the finding that oxaloacetate (OAA), the main physiological substrate of CS, strongly inhibits its thioesterase side-activity (Fig. 1d). Although acetyl-CoA, the other physiological substrate of CS, only reduced its malyl-CoA thioesterase activity by ~30% at the highest concentration tested, oxaloacetate almost completely inhibited the reaction at 50 μM. This suggests that the CS malyl-CoA thioesterase activity only contributes moderately, if at all, to intracellular malyl-CoA degradation.

### TCA cycle enzymes have side-activities producing malyl-CoA

With the literature evidence for enzymatic side-activities forming malyl-CoA^23,24^, commercial porcine heart KGDH was first assayed spectrophotometrically, using KG or KHG as a main and promiscuous substrate, respectively (Fig. 1a, reaction 3)^24^. Indeed, KGDH formed malyl-CoA from KHG and CoA (catalytic efficiency ~16-fold lower than the one for succinyl-CoA formation from KG and CoA) (Table 1). The acyl-CoA forming activities of recombinant human SUCL, produced by co-expressing the *SUCLG1* subunit with the *SUCLA1* subunit (ADP-forming complex) or with the *SUCLG2* subunit (GDP-forming complex) in *E. coli* (Supplementary Fig. 2), were measured spectrophotometrically in the presence of the main substrate (succinate) or the alternative substrates (malate, itaconate) (Fig. 1a, reaction 4)^23^. In agreement with the bacterial homologs^23^, the human SUCL complexes showed higher catalytic efficiencies for itaconyl-CoA formation than for malyl-CoA formation, with catalytic efficiencies that were 30-fold (ADP-forming enzyme) and 15-fold (GDP-forming enzyme) lower with itaconate as with succinate, respectively, while they were 340- and 540-fold lower, respectively, with malate as with succinate (Table 1). Although low, the malyl-CoA forming side-activities measured here with mammalian KGDH and SUCLs are similar or higher, in relative terms compared to the main activities, compared to previously reported metabolic side-activities known to interfere with metabolism^16,17,25^. Our observations thus indicate that promiscuous formation of malyl-CoA is likely to occur in mammalian cells, where it is prone to interfere with normal metabolism if left to accumulate.

**Table 1.** Kinetic characterization of malyl-CoA forming activities of two TCA cycle enzymes. KGDH activities were measured spectrophotometrically at 340 nm (NADH formation) with substrate concentrations varied between 0-2 mM and enzyme concentrations of 88 ng/μL or 175 ng/μL for KG and KHG, respectively. SUCL activities were measured by PK/LDH coupling with a SUCL concentration of 1 ng/μL for the ADP-forming succinyl-CoA ligase activity and of 50 ng/μL for all other activities. Substrate concentrations varied between 0-2 mM succinate, 0-12 mM malate, and 0-50 mM itaconate (ADP-forming complex) or 0-20 mM succinate and 0-150 mM malate or itaconate (GDP-forming complex). Kinetic parameters were estimated in GraphPad Prism (9.5.0) using the non-linear Michaelis-Menten fit model. The values shown represent means ± SD (n=3). Reaction numbers correspond to the Figure 1a reaction scheme. KG, α-ketoglutarate; KHG, 2-keto-4-hydroxyglutarate; KGDH; α-ketoglutarate dehydrogenase complex; SUCL, succinyl-CoA ligase (ADP or GDP-forming complexes).

| Reaction Number | Enzyme | Substrate | $K_M$ (mM) | $V_{max}$ ($\mu\text{mol}\cdot\text{min}^{-1}\cdot\text{mg}^{-1}$) | $k_{cat}$ ($\text{s}^{-1}$) | $k_{cat}/K_M$ ($\text{s}^{-1}\cdot\text{M}^{-1}$) |
| --- | --- | --- | --- | --- | --- | --- |
| 3 | KGDH | KG | $0.22 \pm 0.08$ | $0.04 \pm 0.004$ | $0.07 \pm 0.01$ | $0.32 \pm 0.13 \times 10^3$ |
| 3 | KGDH | KHG | $0.65 \pm 0.32$ | $0.006 \pm 0.001$ | $0.01 \pm 0.002$ | $0.02 \pm 0.01 \times 10^3$ |
| 4 | SUCL<br>(ADP-forming) | Succinate | $6.3 \pm 1.9$ | $4.4 \pm 0.2$ | $6.4 \pm 0.3$ | $1.02 \pm 0.35 \times 10^3$ |
| 4 | SUCL<br>(ADP-forming) | Malate | $21.9 \pm 5.8$ | $0.04 \pm 0.004$ | $0.06 \pm 0.01$ | $0.003 \pm 0.001 \times 10^3$ |
| 4 | SUCL<br>(ADP-forming) | Itaconate | $19.1 \pm 2.5$ | $0.5 \pm 0.6$ | $0.65 \pm 0.08$ | $0.034 \pm 0.002 \times 10^3$ |
| 4 | SUCL<br>(GDP-forming) | Succinate | $0.25 \pm 0.03$ | $9.8 \pm 0.86$ | $13.5 \pm 1$ | $54 \pm 2.1 \times 10^3$ |
| 4 | SUCL<br>(GDP-forming) | Malate | $1.9 \pm 0.3$ | $0.13 \pm 0.01$ | $0.18 \pm 0.02$ | $0.1 \pm 0.01 \times 10^3$ |
| 4 | SUCL<br>(GDP-forming) | Itaconate | $1.5 \pm 0.2$ | $3.8 \pm 0.2$ | $5.3 \pm 0.2$ | $3.5 \pm 0.7 \times 10^3$ |

**Fig. 2.**
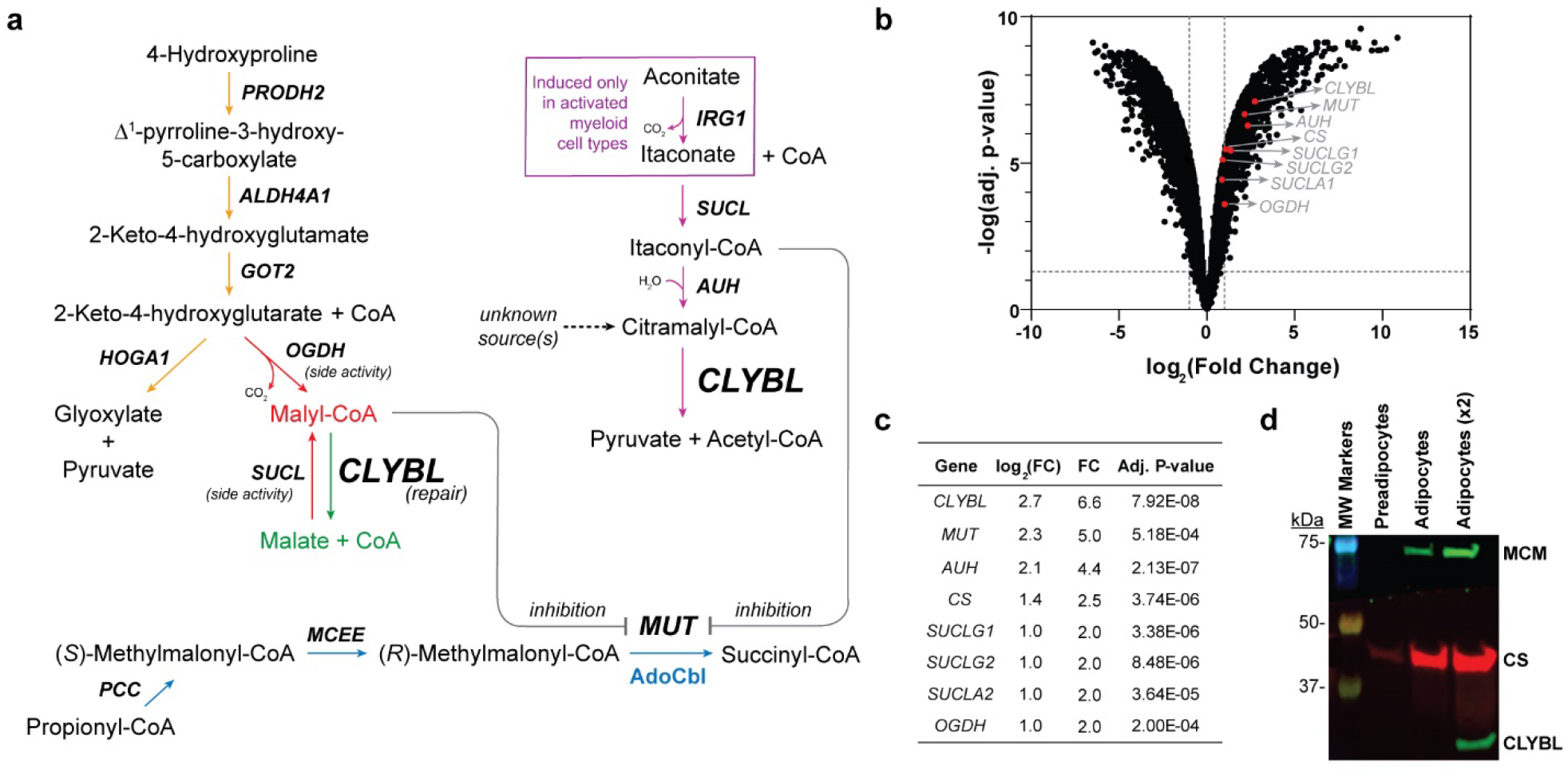
Metabolic roles of CLYBL and enzyme expression in adipocytes. (**a**) Scheme of the known and hypothesized (red: damage formation; green: metabolite repair) metabolic pathways involving CLYBL, including malyl-CoA formation, repair, and MCM inhibition (gray lines). Pathways are also shown for canonical 4-hydroxyproline catabolism (orange arrows), itaconate formation and degradation (purple arrows), and propionyl-CoA catabolism (blue arrows); (**b**) Volcano plot of differential gene expression from deposited microarray data [NCBI Geo: GSE20752] of 3T3-L1 adipogenesis (Day 7 vs. 0) with *CLYBL*-related genes annotated and (**c**) their fold changes and adjusted P-values. (**d**) Western blot analysis of mitochondrial protein extracts of preadipocytes and mature (Day 7) adipocytes (2 μg protein/lane); a 4.5-fold concentrated mature adipocyte extract was loaded in the last lane (9 μg protein). Predicted MWs for the mature proteins (without MTS) for CLYBL, MCM, and CS are 35 kDa, 79 kDa, and 49 kDa, respectively. *CLYBL*, citrate lyase beta-like protein; *PRODH2*, hydroxyproline dehydrogenase; *ALDH4A1*, Δ^1^-pyrroline-5-carboxylate dehydrogenase (mitochondrial); *GOT2*, aspartate aminotransferase (mitochondrial); *HOGA1*, 4-keto-2-hydroxyglutarate aldolase; *OGDH*; α-ketoglutarate dehydrogenase complex component E1; *IRG1*, aconitate decarboxylase; *SUCLG1*, succinyl-CoA ligase (ADP/GDP-forming subunit alpha); *SUCLG2*, succinyl-CoA ligase (GDP-forming subunit beta); *SUCLA2*, succinyl-CoA ligase (ADP-forming subunit beta); *AUH*, 3-methylglutaconyl-CoA hydratase; *PCC*; propionyl-CoA carboxylase; *MCEE*; methylmalonyl-CoA epimerase; *MUT*/MCM (gene/protein), methylmalonyl-CoA mutase; AdoCbl, adenosylcobalamin; *CS*, citrate synthase.

### Malyl-CoA accumulates in *CLYBL* deficient mammalian cells

After identifying plausible metabolic routes for malyl-CoA formation, we knocked out *CLYBL* in HEK293 cells given that the gene is highly expressed in kidney^11^. Failure to detect malyl-CoA in the *CLYBL* KO HEK293 cell lines and our observation, based on public gene and protein expression databases, that C*LYBL* is expressed at low levels in HEK293 cells compared to the brown adipocyte cell line previously used^2^, led us to consider the murine 3T3-L1 adipocyte line for our functional *CLYBL* investigations. To ascertain that our hypothesized metabolite damage and repair pathways involving malyl-CoA (Fig. 2a) are operating in these cells, we analyzed deposited 3T3-L1 microarray gene expression data [NCBI Geo: GSE20752] of preadipocytes (Day 0) and mature adipocytes (Day 7)^26^. All genes potentially involved in malyl-CoA formation (*SUCLG1, SUCLG2, SUCLA2, OGDH*) as well as its degradation (*CLYBL* and *CS*) are expressed in the adipocyte model and, interestingly, were upregulated during adipocyte maturation (Fig. 2b), with the strongest and most significant effect observed for the *CLYBL* gene (Fig. 2c). CLYBL protein could not be detected in mitochondrial extracts of 3T3-L1 preadipocytes or mature adipocytes when loading 2 μg of mitochondrial proteins, but the protein could be detected in the mature adipocytes when using 4.5-fold more concentrated mitochondrial extracts (Fig. 2d). Interestingly, the *MUT* gene, encoding MCM, closely followed *CLYBL* in the list of genes most strongly upregulated during adipogenesis (Fig. 2d), along with CS. Unsurprisingly, expression of *IRG1*, the gene encoding the enzyme responsible for itaconate formation, was not detected in the preadipocytes or mature adipocytes.

We generated two independent *CLYBL* CRISPR knockout lines in 3T3-L1 preadipocytes that were confirmed by Western Blotting analysis (Fig. 3a). Using standard procedures, preadipocytes were differentiated to adipocytes (Extended Data Fig. 1a,b,c,d)^27^ and intracellular metabolites were extracted on Day 12. After solid phase extraction for acyl-CoA enrichment^28^, samples were measured using an optimized untargeted LC-HRMS/MS method. Strikingly, a feature with an *m/z* of 884.1340, and a retention time and MS^2^ spectrum corresponding with the malyl-CoA standard, accumulated in the *CLYBL* KO cells (Fig. 3b,c), in addition to citramalyl-CoA (Fig. 3b,d). Significantly, a 14-fold increase to approximately 0.1 μM malyl-CoA, assuming cells have an average of 200 mg of protein per mL cell volume, was observed in our two independent *CLYBL* KO adipocyte lines compared to control cells (Fig. 3e). These results show, in our knowledge for the first time, that malyl-CoA is formed in mammalian cells and, together with our *in vitro* activity assays described above, suggest that CLYBL acts as a malyl-CoA thioesterase in living cells. Additionally, a 10-fold increase in citramalyl-CoA concentration to approximately 0.2 μM (Fig. 4f) was observed in *CLYBL* KO versus control cells, similar to what was previously observed in brown murine adipocytes^2^ and confirming the citramalyl-CoA lyase activity of CLYBL. Transduction of our *CLYBL* KO lines to express human CLYBL (Extended Data Fig. 1e) restored baseline levels of malyl-CoA and citramalyl-CoA, while their levels remained elevated after transduction with the catalytically null variant hCLYBL^D320A^ (Extended Data Fig. 1f,g), validating that malyl-CoA accumulation is specifically prevented by catalytically active CLYBL. Furthermore, stable isotope labeling confirmed that malate is a direct precursor of malyl-CoA; incubation of *CLYBL* KO cells with 2 mM ^13^C_4_-malate for 48 h led to an 8% molar enrichment in ^13^C_4_-malyl-CoA (Fig. 3g).

**Fig. 3.**
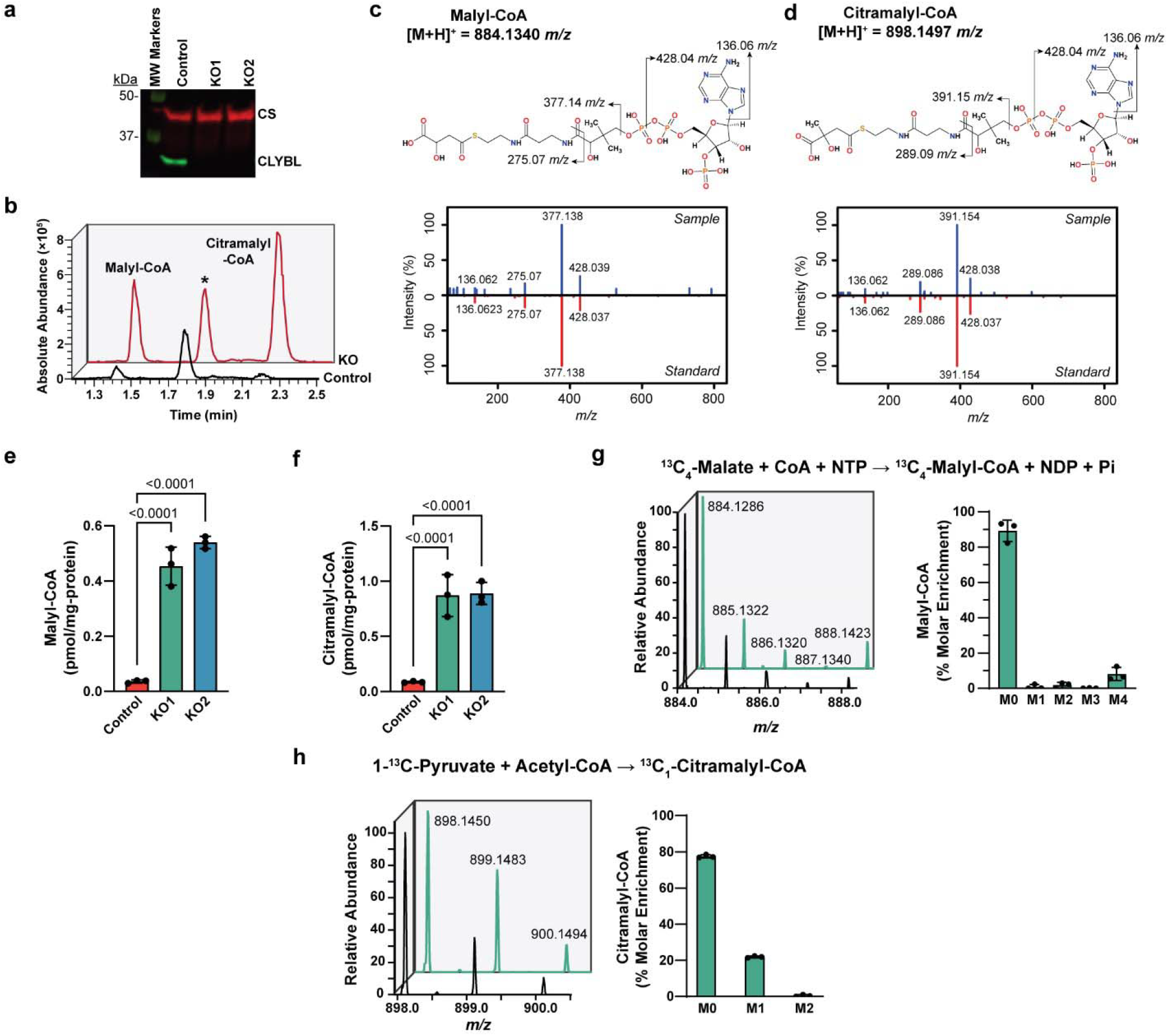
Malyl-CoA and citramalyl-CoA accumulation in *CLYBL* 3T3-L1 KO cells. (**a**) Western blot analysis of mitochondrial fractions (15 μg protein/lane) of mature adipocytes for detection of CLYBL (35 kDa) and CS (49 kDa, loading control) in a control and two polyclonal *CLYBL* knockout (KO1 and KO2) cell lines. (**b**) Representative extracted ion chromatograms (± 10 ppm) of malyl-CoA (calculated [M+H]^+^ = 884.1340 *m/z*) and citramalyl-CoA (calculated [M+H]^+^ = 898.1497 *m/z*; *, unknown feature detected in the same *m/z* window but with dissimilar MS^2^ fragmentation) after LC-HRMS analysis of control and *CLYBL* KO cell extracts. (**c**) Malyl-CoA and (**d**) citramalyl-CoA similarity plots comparing MS^2^ spectra from a *CLYBL* KO cell extract and the standard compound. (**e**) Malyl-CoA and (**f**) citramalyl-CoA concentration in control and *CLYBL* KO cell extracts quantified by LC-HRMS. Stable isotope labeling of *CLYBL* KO1 cells with ^13^C_4_-malate (**g**) or 1-^13^C-pyruvate (**h**). Isotopomer distribution in cell extracts of unlabeled (black) and labeled (green) cells are shown alongside bar charts (right) of the percent molar enrichment of the stable isotope label (^13^C_4_-malyl-CoA, calculated M4 *m/z* = 888.1475; ^13^C_1_-citramalyl-CoA, calculated M1 *m/z* = 899.1531). Data shown represent means ± SD (n = 3), statistical significance was determined through ANOVA followed by Fisher’s LSD *posthoc* test, and the computed p-values are indicated. CLYBL, citrate lyase beta-like protein; CS, citrate synthase.

**Fig. 4.**
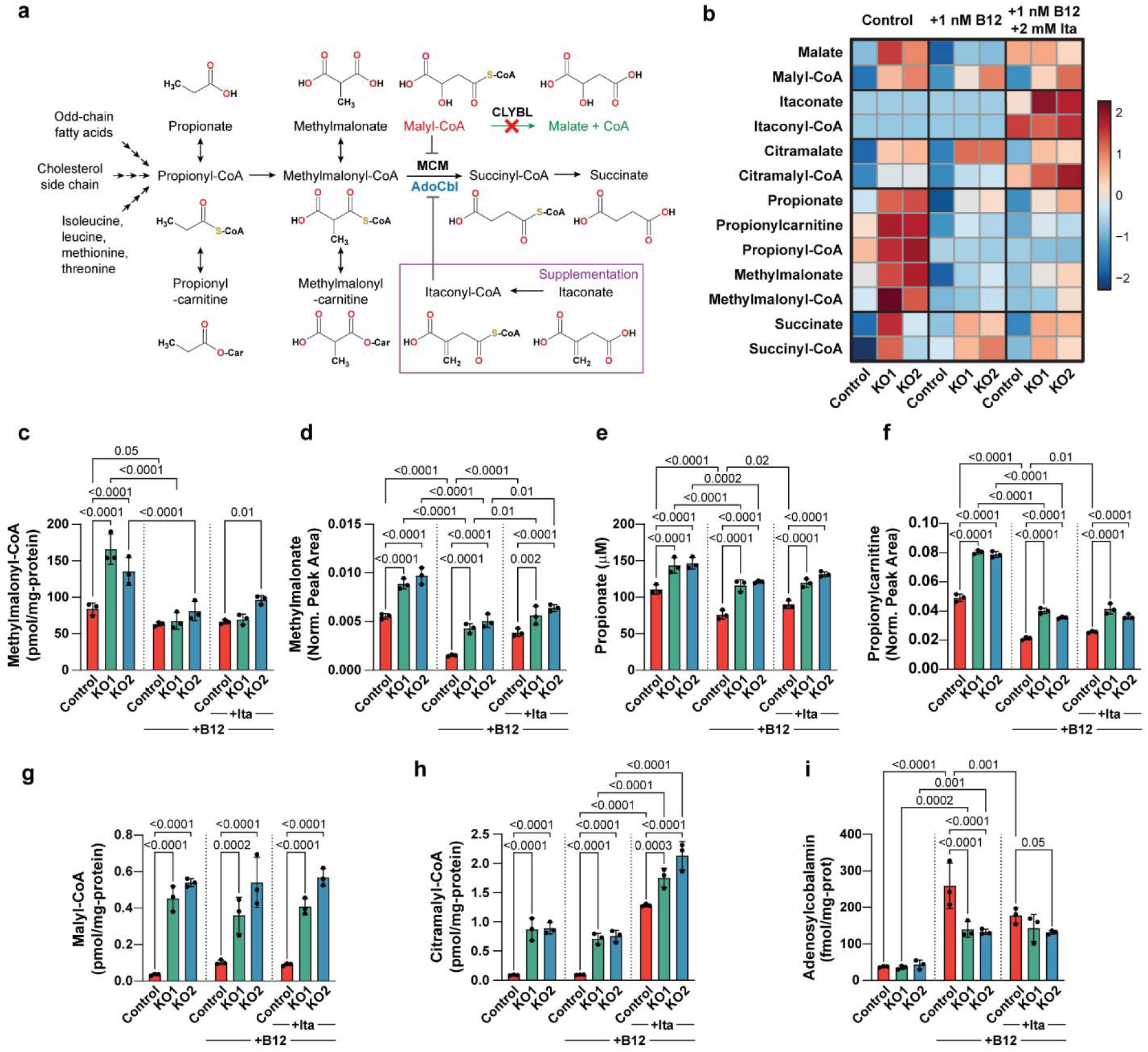
Propionate metabolism perturbation caused by *CLYBL* gene deletion is not exacerbated by itaconate supplementation. The propionate catabolic pathway was profiled in control and *CLYBL* KO adipocytes (Day 12), in the absence or presence of 1 nM B12 (+B12; from Day 0) and/or 2 mM itaconate (+Ita; from Day 4). (**a**) Propionyl-CoA catabolic pathway indicating possible MCM inhibition mechanisms. Propionate and adenosylcobalamin were measured by targeted GC-MS and LC-MS methods, respectively, and all other metabolites by untargeted LC-HRMS/MS methods. (**b**) Heatmap showing autoscaled metabolite level changes in control versus *CLYBL* KO cells, without and with B12 and/or itaconate supplementation. Measured levels of methylmalonyl-CoA (**c**), methylmalonate (**d**), propionate (**e**), propionylcarnitine (**f**), malyl-CoA (**g**), citramalyl-CoA (**h**), and adenosylcobalamin (**i**) in cell extracts, except for propionate (extracellular). Data shown represent means ± SD (n = 3), statistical significance among groups were determined through ANOVA followed by Fisher’s LSD *posthoc* test. Only significant p-values (<0.05) are indicated. Acyl-CoA esters and adenosylcobalamin were quantified using the ion ratio with their surrogate or internal standard (^13^C_3_-malonyl-CoA and ^13^C_3_-caffeine, respectively) and external calibration curves, followed by normalization with total protein. Norm. Peak Area corresponds to data normalized to the surrogate standard (Nε-trifluoroacetyl-L-lysine) only. AdoCbl, adenosylcobalamin; CLYBL, citrate lyase beta-like protein; MCM, methylmalonyl-CoA mutase.

A very intriguing observation, not emphasized previously^2^, is the accumulation of citramalyl-CoA in cells that neither express IRG1 nor were supplied with external itaconate. We postulated that a possible alternative route to citramalyl-CoA formation could involve condensation of pyruvate and acetyl-CoA by an unknown enzyme. Accordingly, incubation of *CLYBL* KO cells with 2 mM 1-^13^C-pyruvate for 48 h resulted in a 22% molar enrichment in ^13^C_1_-citramalyl-CoA (Fig. 3h), substantiating, in our knowledge for the first time, that citramalyl-CoA can form independently of itaconyl-CoA in mammalian cells.

### *CLYBL* deficiency profoundly perturbs propionate metabolism independently of itaconyl-CoA

The previously published data showing a link between CLYBL and B12 depletion^2,7–11^, the mitochondrial co-localization of CLYBL and the B12-dependent enzyme MCM, the structural similarity between malyl-CoA and the MCM product (succinyl-CoA) as well as a known MCM inhibitor (itaconyl-CoA^2^), combined with our findings of intracellular malyl-CoA formation and degradation by CLYBL, led us to hypothesize that MCM inhibition by malyl-CoA may be a cause of decreased B12 levels under *CLYBL* deficiency. Accordingly, measurement of intermediates of the propionyl-CoA pathway, and derivatives thereof (Fig. 4a,b), showed increased levels of intracellular methylmalonyl-CoA, methylmalonate, propionylcarnitine, and extracellular propionate levels in the *CLYBL* KO lines compared to control cells under our standard cultivation conditions (Fig. 4b,c,d,e,f). Propionyl-CoA levels did not significantly change in the *CLYBL* KO versus control lines (Extended Data Fig. 2a) and no carnitine esters could be detected in any of the cell lines for methylmalonate, malate, citramalate, or itaconate. Accumulation of metabolic intermediates upstream of MCM in *CLYBL* KO cells, supported the hypothesis of an inhibitory effect of malyl-CoA on MCM.

As 3T3-L1 cells are effectively in a B12-deficient state under standard cultivation conditions^27,29,30^, we next compared the effect of B12 supplementation. At all concentrations tested (0.5, 1, and 10 nM), B12 reduced propionylcarnitine levels in control cells, but they remained elevated in the *CLYBL* KO versus control lines (Extended Data Fig. 2f). The 1 nM B12 concentration was selected for all subsequent supplementation experiments. This treatment increased intracellular adenosylcobalamin levels by ~7-fold in control cells and led to reduced levels of metabolites upstream of MCM (Fig. 4c,d,e,f; Extended Data Fig. 2a). However, as for propionylcarnitine, methylmalonate and propionate levels remained significantly increased in the *CLYBL* KO versus control cells, with slightly higher fold changes as in the absence of B12 supplementation (Fig. 4d,e). *CLYBL* deficiency did not detectably affect the low adenosylcobalamin levels in untreated cells, but a significant decrease of adenosylcobalamin was measured in *CLYBL* KO versus control cells under B12 replete conditions, indicating an impairment at the level of MCM (Fig. 4i). B12 deficiency under the standard cultivation conditions^27,29,30^ likely causes holo-MCM activity to be near the limit of detection^31^, which may make it difficult to measure the effect of *CLYBL* deficiency (and malyl-CoA accumulation) on adenosylcobalamin levels without B12 supplementation. In standard media, the only (and highly variable) source of B12 is FBS; high, batch-dependent variations in exact composition of the latter may explain the stronger adenosylcobalamin depletion previously observed in *CLYBL* KO brown adipocytes without B12 supplementation^2^. Upon B12 treatment, a 3-fold increase in malyl-CoA levels was observed in the control cells (Fig. 2g), possibly due to increased expression of TCA cycle enzymes with promiscuous activity^32^ and reduced levels of propionyl-CoA (Extended Data Fig. 2a), a TCA cycle inhibitor^33^.

Since itaconate should not be produced by adipocytes under our standard cultivation conditions, we tested the effect of external itaconate supplementation (at 2 mM) on the propionyl-CoA pathway in our adipocyte cell lines. Itaconate supplementation was previously shown by Shen *et al*.^2^ to lead to citramalyl-CoA accumulation in wildtype brown adipocytes and other cell types, but *CLYBL* KO cells were not challenged with itaconate supplementation in this study. We predicted itaconate supplementation to have a more severe impact on control cells than on *CLYBL* KO cells where the propionate pathway was already inhibited by malyl-CoA. As expected, itaconate and itaconyl-CoA, were only detected in cells supplemented with itaconate (Extended Data Fig. 2b,c). A more than 10-fold increase in citramalyl-CoA levels was also observed in control cells upon itaconate supplementation and these levels further increased in the *CLYBL* KO lines (Fig. 4h). However, the KO to WT ratio for citramalyl-CoA levels was much higher in untreated (~10-fold) than in itaconate supplemented conditions (1.5-fold), whereas for malyl-CoA levels this ratio was similar in control and itaconate supplemented conditions (Fig. 4g). As predicted, itaconate supplementation had a more pronounced effect on metabolites upstream of MCM (methylmalonate, propionate, and propionylcarnitine) in control cells than in *CLYBL* KO cells; in fact, itaconate supplementation did not significantly change the levels of these metabolites in the KO cells (Fig. 4d,e,f). Similarly, itaconate supplementation led to a 1.5-fold decrease in adenosylcobalamin levels in control cells (Fig. 4i) but had no effect on the cofactor levels in the *CLYBL* KO cells, where MCM was presumably already maximally inhibited by malyl-CoA. Taken together, our results show that itaconate supplementation did not exacerbate MCM inhibition in the *CLYBL* KO cells and, importantly, that this inhibition can occur completely independently of itaconyl-CoA, likely via the CLYBL substrate malyl-CoA.

### Malyl-CoA inhibits methylmalonyl-CoA mutase via coenzyme B12 inactivation

As our findings suggested an itaconate-independent MCM inhibition in *CLYBL* deficient adipocytes, we sought for direct evidence of MCM inhibition by malyl-CoA. First, we tested the effect of malyl-CoA or itaconyl-CoA on the enzymatic activity, by preincubating recombinant human MCM (Supplementary Fig. 3) for 10 min with either acyl-CoA before addition of the methylmalonyl-CoA substrate at a concentration corresponding to the reported *K*_m_ (65 μM^12^). Both acyl-CoAs induced a dose-dependent inhibition of MCM (Fig. 5a), with malyl-CoA exerting a more potent effect than itaconyl-CoA. MCM activity was reduced to <10% at the highest malyl-CoA concentration (50 μM) tested (control assays with free CoASH, contained as impurity in our synthesized itaconyl-CoA standard (Supplementary Fig. 6c) did not affect MCM activity). Next, we examined whether malyl-CoA stabilizes the formation of MCM-bound cob(II)alamin, an intermediate formed during the MCM catalytic cycle. Expectedly, the addition of methylmalonyl-CoA to MCM preincubated with adenosylcobalamin had no effect on the absorbance spectrum (Fig. 5b). However, the addition of malyl-CoA induced a rapid absorbance shift of λ_max_ from 530 nm to 466 nm (Fig. 5c), indicating a conversion of the MCM-bound adenosylcobalamin into cob(II)alamin. We observed the same change in absorbance spectrum upon addition of itaconyl-CoA (Fig. 5d), as was also reported previously^2,12^. Given that malyl-CoA should be more ubiquitously formed across tissues than itaconyl-CoA and is a more potent inhibitor of MCM than the latter, accumulation of this non-canonical acyl-CoA when *CLYBL* is deficient may represent the main mechanism underlying reduced levels of circulating vitamin B12 in individuals lacking this repair enzyme activity.

**Fig. 5.**
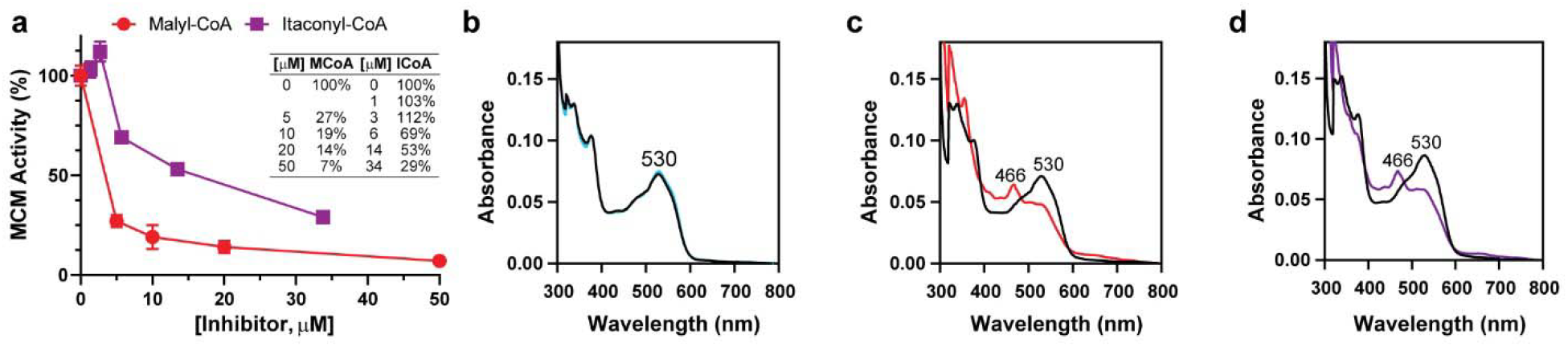
Malyl-CoA causes methylmalonyl-CoA mutase inhibition and coenzyme depletion. (**a**) MCM activity in the presence of increasing concentrations of malyl-CoA (red circles) or itaconyl-CoA (purple squares). Relative activities compared to the activity in absence of inhibitor (100%) are shown. MCM (0.12 μM) was incubated for 10 min with 0.24 μM adenosylcobalamin and inhibitor at the indicated concentrations. The methylmalonyl-CoA substrate (65 µM) was then added, and reactions were quenched 10 min later with 10% formic acid to measure succinyl-CoA levels by LC-HRMS/MS. The values shown are means ± SD (n = 3). Representative absorbance spectra of MCM (12.5 µM, homodimer) preloaded with 5 μM adenosylcobalamin (black traces) and after a 25-min incubation with methylmalonyl-CoA (blue trace, **b**), malyl-CoA (red trace, **c**), or itaconyl-CoA (purple trace, **d**). The shift in the λ_max_ from 530 nm to 466 nm indicates the conversion of adenosylcobalamin to cob(lI)alamin in the active site of MCM. MCM, methylmalonyl-CoA mutase.

## Discussion

Biology is inherently imperfect, and underground networks of enzymatic repair reactions exist to eradicate toxic or useless metabolites formed by enzymatic side-activities and unwanted chemical reactions^14,15^. Herein, we report the identification of an additional, more ubiquitous metabolic role for CLYBL than its function in itaconate catabolism, that is more compatible with its wide tissue distribution, namely, as a metabolite repair enzyme for the non-canonical metabolite malyl-CoA to avert inhibition of MCM. Given the reported citramalyl-CoA lyase activity of CLYBL, we expected CLYBL to similarly convert malyl-CoA to acetyl-CoA and glyoxylate, but instead found that CLYBL acts as a thioesterase specific for malyl-CoA. CLYBL thus seems to harbor two types of enzymatic activities in the same catalytic site. We found that the catalytic efficiency of the previously uncharacterized malyl-CoA thioesterase activity of CLYBL is 12-fold greater compared to its citramalyl-CoA lyase activity. We confirmed that CS also acts as a malyl-CoA thioesterase *in vitro*^24^, albeit with a 5-fold lower catalytic efficiency compared to CLYBL. In addition, one of the main substrates of CS, oxaloacetate, strongly inhibited this thioesterase side-activity of CS at physiological concentrations (2-5 μM in mitochondria^34^). During the physiological reaction catalyzed by CS, oxaloacetate binds first to CS, which increases the binding constant for the second substrate, acetyl-CoA^35^. The latter only poorly inhibited the malyl-CoA thioesterase activity of CS. Taken together, these results suggest that CS does not substantially contribute to malyl-CoA degradation *in vivo*, except perhaps under conditions of oxaloacetate depletion. The accumulation of malyl-CoA in the *CLYBL* KO lines shows, in our knowledge for the first time, that this non-canonical metabolite can form in mammalian cells and that CLYBL plays a main role in its degradation. Furthermore, we showed that C1-labeled pyruvate is incorporated into citramalyl-CoA that accumulated in *CLYBL* KO cells in the absence of itaconate supplementation, providing first evidence for the existence of an alternative pathway to citramalyl-CoA formation in mammalian cells.

Due to the high flux through certain metabolic pathways, and potentially high concentrations of substrate analogs feeding into enzymatic side-activities, even extremely low side-activities can lead to the formation of side products at levels that perturb cellular metabolism if left to accumulate (*i*.*e*., in the absence of a metabolite repair activity). A striking example is the formation of L-2-hydroxyglutarate, a side product of malate dehydrogenase, whose accumulation leads to a severe neurometabolic disease caused by a repair enzyme deficiency; L-2-hydroxyglutarate is formed with a 10^7^-fold lower efficiency by malate dehydrogenase (from α-ketoglutarate) as compared to its main product malate (from oxaloacetate)^36^. Furthermore, glyceraldehyde-3-phosphate dehydrogenase converts erythose-4-phosphate to the non-canonical 4-phosphoerythronate at a 3500-fold lower efficiency compared to the efficiency with its main substrate^25^. This minor side activity leads to an accumulation of 20-30 μM 4-phosphoerythronate in cells deficient in the repair enzyme phosphoglycolate phosphatase, interfering with both the pentose phosphate pathway and glycolysis^25^. Two enzymes had previously been reported to form malyl-CoA *in vitro* via promiscuous activities: bacterial SUCL^23^ (when using malate instead of succinate) and porcine heart KGDH^24^ (when using KHG instead of KG). We determined catalytic efficiencies for the malyl-CoA producing side-activity that were ‘only’ 340-, 540-, and 16-fold lower compared to the ones of the main activities of SUCL (ADP-forming, GDP-forming) and KGDH, respectively. This strongly suggests that TCA cycle enzymes form malyl-CoA as a side product in cells at levels that are prone to interfere with other metabolic activities. SUCL, notably, is ubiquitously expressed in mitochondria where it is exposed to high concentrations of the promiscuous substrate, malate, which we identified as a direct precursor of malyl-CoA with stable isotope labeling. Normally degraded by KHG aldolase^24^, KHG, the promiscuous substrate of KGDH, is a catabolite of 4-hydroxyproline that is primarily degraded in the kidney and liver^37^, which are among the tissues where CLYBL is most highly expressed^11^. CLYBL is also highly expressed in skeletal muscle and heart^11^, which are rich in connective tissue, in addition to adipose tissue, in which collagen-related genes are upregulated early in adipogenesis^38^.

*CLYBL* deficiency has been strongly associated through GWAS with decreased levels of circulating vitamin B12^7–11^. The underlying mechanism most likely involves an inhibitory interaction of acyl-CoA esters normally degraded by CLYBL with the mitochondrial B12-dependent enzyme MCM, hallmarked by an accumulation of propionate metabolites (Shen *et al*., (2017)^2^ and this study) which emulates some aspects of inborn errors of propionate and B12 metabolism^39,40^. The upregulation of CLYBL and MCM during adipogenesis supports a role for CLYBL in maintaining functional propionate metabolism. Supplementation of our cell culture media with B12 led to decreased levels of metabolites upstream of MCM in both the control and *CLYBL* KO cells compared to the untreated condition, agreeing with previous studies showing that 3T3-L1 cells are effectively deficient in B12 and hence, propionyl-CoA metabolism under standard cultivation conditions^27,29,30^. This observation also suggests that B12 supplementation in *CLYBL* deficient individuals may counteract adverse effects of methylmalonate and propionate accumulation. However, we continued to observe hallmarks of MCM inhibition in our *CLYBL* KO cells compared with control cells under B12 repletion, including a significant decrease of adenosylcobalamin, showing that B12 homeostasis remained impaired under CLYBL deficiency, presumably through loss of the cofactor at the level of MCM. A radical transfer reaction between the MCM-bound 5’-deoxyadenosylcobalamin and the methylene group of itaconyl-CoA leads to the formation of cob(II)alamin and an adduct which prevents cofactor repair by cobalamin adenosyltransferase^12^. These observations provided leads towards understanding the adenosylcobalamin depletion observed in control cells (adipocytes, HEK293T, and human B-lymphocytes) supplemented with itaconate or RAW264.7 macrophages stimulated with LPS^2^. The depletion of adenosylcobalamin in macrophages was confirmed to be dependent on *IRG1* induction^12^, uncovering a mechanism of MCM inhibition (via itaconyl-CoA) in activated immune cell types and in engulfed pathogens. The same phenotype of adenosylcobalamin depletion was observed in *CLYBL* KO adipocytes in the absence of itaconate supplementation (this study and Shen *et al*.^2^), where neither itaconate nor itaconyl-CoA could be detected (this study). *IRG1* expression is inducible and highly cell-type specific and was not detected in the 3T3-L1 gene expression data, confirming the generally accepted notion that itaconate production is restricted to immune cell types. Widely distributed enzymes (SUCL, 3-methylglutaconyl-CoA hydratase (AUH), CLYBL)^1,41,42^ are responsible for itaconate catabolism, and thus itaconyl-CoA production via SUCL can be elicited in many cell types when supplemented with itaconate. The latter treatment altered branched-chain amino acid catabolism and increased odd-chain fatty acid levels in liver cells via activation to itaconyl-CoA and inhibition of MCM^43^. Itaconate supplementation in our B12-repleted 3T3-L1 control adipocytes led to less pronounced effects on propionate catabolism than *CLYBL* gene deletion and itaconate treatment did not exacerbate the effects on propionate catabolism in *CLYBL* KO cells. These results suggest that the substrate for CLYBL that we identified in this study, malyl-CoA, perturbs propionate metabolism via a similar mechanism than itaconyl-CoA.

It remains unclear why *CLYBL* deficiency is seemingly well tolerated in the human population (the LOF SNP rs41281112 is present in 2.7% of human chromosomes^6^), although it leads to a secondary vitamin B12 deficiency and possible consequences, including an increased risk of neurodegeneration, especially in subjects with diets low in vitamin B12 and/or impaired B12 absorption^44^. One evolutionary advantage that may actually be conferred by *CLYBL* deficiency is a more efficient clearance of pathogens engulfed by macrophages, due to increased levels of the bactericidal itaconyl-CoA as a consequence of the inability to catabolize citramalyl-CoA^12^. However, we did not observe significant differences in itaconyl-CoA levels between our control and *CLYBL* KO cell lines (after supplementation with itaconate), indicating that metabolite accumulation upstream of *CLYBL* blockage may be limited to the most proximal intermediates (citramalyl-CoA and maybe mesaconyl-CoA). Many pathogens are dependent on B12 from their hosts^45^; thus, as an alternative explanation, *CLYBL* deficiency may decrease susceptibility to certain infectious agents by depriving them of vitamin B12^2,12^.

Taken together, our cell culture results suggest a significant role for malyl-CoA as a mediator between CLYBL and B12 deficiency and this is further supported by our observation that malyl-CoA inhibited the enzymatic activity of MCM more effectively than itaconyl-CoA. We further provide evidence that malyl-CoA renders MCM inactive by stabilizing its catalytic intermediate cob(II)alamin, elucidating the mechanism of the observed adenosylcobalamin depletion. Although cob(II)alamin formation is not necessarily an indicator of a 5’-deoxyadenosyl radical transfer mechanism^46^, malyl-CoA is conceivably a suicide inhibitor and seems to prevent damaged cofactor off-loading by the MCM repair machinery as shown previously for itaconyl-CoA^12^. The intrinsic reactivity of cofactors and coenzymes cause them to be frequent targets, or initiators, of enzymatic or spontaneous damage^14^. We are just starting to unveil messy primary metabolism, and many damaged versions of cofactors and coenzymes may remain unknown, in addition to the repair enzymes responsible for maintaining them under their forms which are essential for life.

## Methods

### Enzymatic synthesis and purification of malyl-CoA, citramalyl-CoA, and itaconyl-CoA

Recombinant *Chloroflexus aurantiacus* malyl-CoA/citramalyl-CoA lyase (CaMcl) containing an N-terminal hexahistidine tag was generated for malyl-CoA/citramalyl-CoA production, similarly to previously described methods^1^. Gateway cloning was used to prepare the bacterial expression plasmid by inserting the CaMcl coding sequence (GenBank ID AGR55786.1) into a pDONR221 entry vector (Invitrogen) via a BP reaction, following the manufacturer’s instructions. The insert was transferred to a pDEST527 vector (Addgene plasmid #11518) using an LR clonase reaction. The resulting construct was confirmed by sequencing and transformed into *E. coli* RosettaBlue(DE3) cells for protein overexpression. Overnight precultures were grown at 37 °C in Luria-Bertani (LB) media containing 1% glucose and 100 μg/mL ampicillin and used to inoculate main cultures in the same media but without glucose. Protein overexpression was induced at an OD_600_ of 0.5, after cold-shocking the cells on ice for 30 min, by addition of 0.5 mM isopropylthiogalactopyranoside (IPTG) and overnight cultivation at 22 °C. Cells were harvested 16-18 h post-induction by centrifugation for 15 min at 4 °C and 4500 *g*. Cells were lysed by freeze-thawing in a buffer containing 20 mM HEPES (pH 7), 1 mM dithiothreitol (DTT), 10 mg/mL lysozyme, and cOmplete Protease Inhibitor (Roche, 05892791001, 4693132001). The cell lysate was incubated for 30 min at 4 °C with 0.1 mg/mL DNAse I from bovine pancreas (Roche 11284932001) and 10 mM MgSO_4_, followed by centrifugation for 40 min, 4 °C at 17000 *g* and filtration of the supernatant (0.45 μm filter). Protein purification was conducted on a nickel affinity column (HisTrap HP, 1mL, GE Healthcare, 17-5247-01) using an ÄKTA Pure 25M Chromatography System (GE Healthcare) with buffers (A) 25 mM Tris (pH 7.5), 300 mM NaCl, and 10 mM imidazole and (B) 25 mM Tris (pH 7.5), 300 mM NaCl, and 500 mM imidazole. After loading the filtered cell extract, the column was washed with buffer A, followed by a second wash step at 3% buffer B for 10 mL, prior to applying a linear gradient of 3% - 100% buffer B over 20 min. Peak fractions were pooled and desalted using a HiTrap Desalting column (5 mL, Cytiva, 17140801) with a buffer containing 25 mM Tris (pH 7.5) and 25 mM NaCl. Desalted fractions were pooled and supplemented with 10% glycerol before storage at −80 °C. The final purified protein preparation was analyzed by SDS-PAGE and a >95% purity was estimated using Coomassie staining (Supplementary Fig. 4).

A pET30a plasmid containing the coding sequence for *Pseudomonas aeruginosa* succinyl-CoA:itaconate CoA transferase (PaIct) with a hexahistidine tag was purchased from Addgene (plasmid #111293)^2^ and transformed into *E. coli* BL21 cells for overexpression. Following established methods^1^, precultures were grown overnight in LB containing 50 μg/mL kanamycin at 37 °C and shaking at 200 rpm and used to inoculate the main culture in the same medium. Once the OD_600_ reached 1.2, the culture was cold-shocked on ice for 30 min and 0.5 mM IPTG was added. Protein overexpression was performed overnight at 18 °C with shaking at 200 rpm. Cells were harvested by centrifugation for 15 min at 4 °C and 4500 *g* and the cell pellets were stored at −80 °C until protein purification. Protein purification was performed as described above for CaMcl, except that cells were lysed by tip sonication for 2 min with 0.5 sec pulses (50% amplitude) every 2.5 sec in an extraction buffer containing 25 mM Tris (pH 8), 500 mM NaCl, 0.5 mM phenylmethylsulfonyl fluoride (PMSF), and 1 mM DTT. The cell lysate (not DNase treated) was centrifuged for 30 min at 4 °C and 17000 *g*, the supernatant was filtered on a 0.45 μm cellulose acetate filter and finally supplemented with 10 mM imidazole. The eluents used for protein purification were: (A) 25 mM Tris (pH 8), 500 mM NaCl, and 10 mM imidazole and (B) 25 mM Tris (pH 8), 500 mM NaCl, and 300 mM imidazole. Desalted fractions were pooled, supplemented with 10% glycerol and 100 μg/mL BSA, and stored at −80 °C until further use.

Previously described methods were used to synthesize malyl-CoA/citramalyl-CoA^1^ and itaconyl-CoA^2^, except 250 mM Tris (pH 7.5) was used as the buffer. Purification of acyl-CoA esters was conducted using a Shimadzu UHPLC Nexera X2 equipped with a PDA detector and a semi-preparative Luna 5u C18(2) (250 × 10 mm, 100Å; Phenomenex, 00G-4252-NO) column. The chromatography was performed by applying mobile phases (A) 40 mM ammonium formate (pH 4.5) and (B) acetonitrile with a flow rate of 4 mL/min and the following gradient: 0 min, 4% B; 5 min, 4% B; 10 min, 7% B; 25 min, 9% B; 30 min, 12% B; 33 min, 12% B; 35 min, 4% B; 45 min, 4% B. Acyl-CoA peaks were detected at 260 nm and 1 mL fractions were collected (Supplementary Fig. 5). Peak fractions were pooled, aliquoted, and dried by speedvac overnight at 4 °C followed by 20 min at 20 °C before storage at −20 °C. Aqueous solutions prepared by resuspension of acyl-CoA esters were titrated at 260 nm using the extinction coefficient *ε* = 16400 M^−1^cm^−1^. Identity of the purified standards was confirmed using high-resolution LC-MS (MS^1^ and MS^2^). Malyl-CoA, citramalyl-CoA, and itaconyl-CoA had an estimated purity of 97%, 96%, and 27%, respectively, based on HPLC detection at 260 nm (Supplementary Fig. 6). Dried itaconyl-CoA stocks were reconstituted immediately before use to minimize degradation.

### Malyl-CoA thioesterase and citramalyl-CoA lyase activity measurements

The full-length ORF of the *CLYBL* coding sequence (GenBank ID AAH34360.1) was PCR amplified from human skin (squamous cell carcinoma) cDNA and ligated into the HindIII (5’-CATC**AAGCTT**ATGGCGCTACGTCTGCTGC-3’) and XhoI (5’-CTTG**CTCGAG**TTTTTCCTTGATGGAGGTGGC-3’) sites (indicated in bold) of the pCMV6-Entry Vector (OriGene, PS100001). The resulting overexpression plasmid, fusing a C-terminal Myc-DKK (*i*.*e*., FLAG) tag to CLYBL, was shown by sequencing to contain the coding sequence of the *CLYBL* natural variant VAR_032101 241 (dbSNP:rs3783185) with an Ile241Val mutation compared to the reported canonical sequence (UniProt Q8N0×4). HEK293FT cells maintained in DMEM (Gibco, 11995065) culture medium containing 4.5 g/L glucose, 1 mM pyruvate, 4 mM glutamate, 1% penicillin/streptomycin, and 10% FBS at 37 °C with 5% CO_2_ were used for protein overexpression. Cells were seeded in 10 cm^2^ Petri dishes (1×10^6^ per dish) and transfected the next day by addition of a complex containing 8 μg of the overexpression plasmid and 16 μL of JetPEI (Polyplus, 101-10N) per dish. Cultures were stopped 48 h later, after aspiration of the medium and scraping of the cells into a lysis buffer (4 mL per dish) containing 50 mM Tris-HCl (pH 7.5) and cOmplete ULTRA EDTA-free protease inhibitor cocktail (Roche). Lysates were transferred to a 15 ml Falcon tubes, flash-frozen in liquid nitrogen, and stored at −80 °C. Protein extracts were prepared by thawing the lysates on ice and DNase treatment as described above. Crude extracts were centrifuged for 30 min at 16,400 *g* and 4 °C and NaCl was added to the supernatants at a final concentration of 150 mM. Recombinant CLYBL was purified under gravity flow on columns packed with 200 μL ANTI-FLAG M2 Affinity Gel (Sigma-Aldrich, A2220) according to the manufacturer’s instructions. Protein extracts (4 mL) were loaded onto the resin equilibrated with 10 column volumes of Tris buffer saline (TBS) solution. Columns were washed with 2.5 mL TBS prior to elution of bound proteins with 5 × 200 μL of 0.1 mg/mL FLAG peptide (Sigma-Aldrich, F4799) in TBS. The first elution fraction was discarded, and the remaining elution fractions (containing CLYBL) were pooled and stored at −80 °C after the addition of glycerol to a final concentration of 10%. Purity of the resulting CLYBL preparation was estimated at >95% by SDS-PAGE and Coomassie staining (Supplementary Fig. 1).

CLYBL thioesterase activity was assayed spectrophotometrically in a Tecan plate reader at 412 nm and 37 °C, using Ellman’s reagent (5,5-dithio-bis-(2-nitrobenzoic acid); Sigma-Aldrich D8130). The assays were performed in a mixture (100 μL total volume) containing 50 mM HEPES (pH 7.5), 2 mM MgCl_2_, 0.1 mM Ellman’s reagent, and the acyl-CoA substrate at the indicated concentration and the reaction was launched by addition of 10 ng purified recombinant CLYBL. For the substrate specificity assay, acyl-CoAs were added at a final concentration of 100 μM (acetyl-CoA, A2056; butyryl-CoA, B1508; DL-3-hydroxybutyryl-CoA, H0261; DL-3-hydroxy-3-methylglutaryl-CoA, H6132; malonyl-CoA, M4263; methylmalonyl-CoA, M1762; n-propionyl-CoA, P5397; and succinyl-CoA, S1129 were from Sigma-Aldrich; acetoacetyl-CoA, SC-252348, was from Santa Cruz Biotechnology). For determination of the kinetic parameters of the malyl-CoA thioesterase activity of CLYBL, malyl-CoA was added at final concentrations ranging from 0-90 μM. Citrate synthase (CS) thioesterase activity was measured similarly at 37 °C in 100 μL reactions containing 100 mM Tris (pH 7.5), with 0.2 mM Ellman’s reagent, 0-700 μM malyl-CoA, and 300 ng porcine heart CS (Sigma-Aldrich, C3260). CS malyl-CoA thioesterase inhibition experiments were conducted using the same assay, with a malyl-CoA concentration of 100 μM (near K_*M*_ value) and acetyl-CoA or oxaloacetate concentrations ranging from 0-100 μM. All assays were background corrected for control reactions where acyl-CoA was added, except for the succinyl-CoA thioesterase activity of CLYBL where the observed slope during the 3-5 min of equilibration time prior to the addition of enzyme was first subtracted to correct for spontaneous succinyl-CoA degradation.

Citramalyl-CoA lyase activity was measured in a coupled reaction with lactate dehydrogenase (LDH) and the consumption of NADH was measured by monitoring A_340_ at 37 °C. The 100 μL assay mixture contained 50 mM HEPES (pH 7.5), 0-300 μM citramalyl-CoA, 2 mM MgCl_2_, 0.5 mM NADH (Roche, 10107735001), 3 U/mL LDH (Sigma-Aldrich, 9001-60-9), and 294 ng CLYBL (added to launch the reaction). All assays were background corrected for control reactions where citramalyl-CoA was not added.

### Measurement of malyl-CoA forming side-activities of TCA cycle enzymes

Porcine heart α-ketoglutarate dehydrogenase (Sigma-Aldrich, K1877) was used to compare the enzymatic activity of this enzyme on α-ketoglutarate versus D,L-2-keto-4-hydroxyglutarate (Sigma-Aldrich, 96599) by spectrophotometric measurement of NADH formation (A_340_) at 30 °C^24^. The 200 μL reaction mixture consisted of 50 mM potassium phosphate (pH 7.5), 1 mM MgCl_2_, 0.33 mM NAD^+^, 0.05 mM coenzyme A, 0.1 mM thiamine phosphate, 3 mM cysteine, and 0-2 mM of the dicarboxylic acid substrate. To launch the reactions, 17.6 μg and 35.0 μg of KGDH were used in the presence of α-ketoglutarate and D,L-2-keto-4-hydroxyglutarate, respectively. Control assays without α-ketoglutarate or D,L-2-keto-4-hydroxyglutarate added to the reaction mixture was used for background correction.

To measure a putative malyl-CoA forming activity of succinyl-CoA ligase, expression vectors containing the ORFs of *SUCLG1* (NCBI RefSeq NP_003840.2; corresponding to amino acids 28-333) fused to an N-terminal hexahistidine tag (pET28a vector) and of *SUCLG2* (NCBI RefSeq NP_003841.1; corresponding to amino acids 38-420) and *SUCLA1* (NCBI RefSeq NP_003839.2; corresponding to amino acids 53-463) fused to a C-terminal hexahistidine tag (two separate pET31a vectors) were purchased from Genscript (all inserts were codon optimized for expression in *E. coli*). Plasmids containing (1) *SUCLG1* (ADP/GDP-forming subunit alpha) and *SUCLA2* (ADP-forming subunit beta) and (2) *SUCLG1* and *SUCLG2* (GDP-forming subunit beta) were co-transformed into competent *E. coli* BL21 cells and positive clones were selected on medium containing ampicillin and kanamycin. Single clones were inoculated in LB containing 50 μg/mL kanamycin and grown at 37 °C overnight. Overnight precultures obtained from single recombinant clones were used to inoculate the main culture in Super Broth containing 50 μg/ml kanamycin and 100 μg/ml ampicillin, which was grown until an OD_600_ of 0.4-0.8 was reached. The cultures were cold-shocked on ice for 30 min and protein expression was induced by addition of 0.4 mM IPTG followed by 40-48 hours incubation at 16 °C and shaking at 200 rpm. Cells were harvested by centrifugation at 4,200 *g* for 20 min (4 °C). Cell pellets were resuspended in lysis buffer containing 50 mM Tris-HCl (pH 7.8), 1 M NaCl, 75 mM imidazole, 1 mg/mL of lysozyme, and stored at −80 °C until purification. Samples were thawed at room temperature and cells were lysed on ice by tip sonication (as described above). Samples were centrifugated at 23,000 *g* for 30 min (4 °C), and the supernatant was filtered (0.45 μm filter) and loaded on a 5 mL HisTrap Fast Flow column (GE Healthcare, 17-5255-01). The purification was conducted on the bench using a peristaltic pump. After washing the column with 25 mL of 50 mM Tris-HCl (pH 7.6), 1 M NaCl, and 75 mM imidazole, His-tagged proteins were eluted with 50 mM Tris-HCl (pH 7.5), 150 mM NaCl, and 200 mM imidazole. Active fractions were collected, concentrated on an Amicon filter with a 10,000 Da MW cut-off, and the buffer was exchanged to 50 mM Tris-HCl (pH 7.2) containing 100 mM NaCl and 10% glycerol. Concentrated purified proteins (1-2 mg/mL) were stored at −80 °C until use. Protein purity was estimated to >95% by SDS-PAGE analysis with Coomassie staining (Supplementary Fig. 2).

SUCL (ADP- and GDP-forming complexes) enzymatic activities were measured spectrophotometrically at 30 °C by coupling the formation of ADP or GDP with pyruvate kinase (PK) and LDH, as previously described^23^. The reaction mixture (100 μL total volume) consisted of 50 mM Tris (pH 7.5), 7 mM MgCl_2_, 7 mM KCl, 0.2 mM NADH (Roche, 10107735001), 0.1 mM CoA (Sigma-Aldrich, C3144), 2 mM phosphoenolpyruvate (Sigma-Aldrich, P7127), 0.6 mM ATP or GTP, 60 U/mL LDH (Sigma-Aldrich, 9001-60-9), and 60 U/mL PK (Sigma-Aldrich, P9136-5KU). For measuring the activity of the ADP-forming enzyme (SUCLG1:SUCLA2 complex) with succinate as a substrate, 0.1 μg of enzyme was added to the 100 μL reaction mixture, while 5 μg of the ADP-or GDP-forming complex was used for all other enzymatic reactions tested. For the GDP-forming reactions (SUCLG1:SUCLG2 complex), tested substrate concentration ranges were: 0-2 mM succinate, 0-12 mM malate, and 0-50 mM itaconate, while substrate concentrations for the ADP-forming reactions (SUCLG1:SUCLA2 complex) ranged from 0-20 mM succinate and 0-150 mM malate or itaconate. Reactions containing no organic acid substrate were used for background corrections.

### Methylmalonyl-CoA mutase (MCM) inhibition assays

A pET28b(+) vector containing a codon optimized full-length ORF of human *MUT* (NCBI RefSeq NP_000246.2) fused to an N-terminal hexahistidine tag was purchased from GenScript^47,48^. Site-directed mutagenesis with the QuickChange Lightning kit (Agilent, 210518-5) was used to remove the mitochondrial targeting sequence (amino acids 1-33) from the vector, following the manufacturer’s instructions and primers designed using the QuickChange Primer Design Program: 3’-ggggctgttgctggtgcatccatggtatatct-5’ and 5’-agatataccatggatgcaccagcaacagcccc-3’. Correct mutagenesis was confirmed by Sanger sequencing using T7 forward and reverse primers. The plasmid was transformed into *E. coli* BL21 cells and the latter were cultured in terrific broth at 37 °C until OD_600_ reached 1-2^49^. After 30 min cold-shock on ice, cultures were induced with 1 mM IPTG, supplemented with 3% DMSO to improve protein stability^47^, and shaken at 200 rpm overnight at 18 °C. A total volume of 6 L of culture were collected per purification. Cell pellets were harvested by centrifugation, as described above, and stored at −80 °C. Cells were resuspended in a lysis buffer containing 50 mM Tris (pH 8), 500 mM NaCl, 20 mM imidazole, 0.5 mM PMSF, 1 mM DTT, 5% glycerol, 1 μL per 10 mL of DNase 1 (Sigma-Aldrich, D5307), 1 mM MgSO_4_, and cOmplete ULTRA EDTA-free protease inhibitor cocktail (Roche) and lysed by tip sonication for 4 min at an amplitude of 40% with 15 s pulses separated by 60 s breaks. Lysates were centrifuged and filtered prior to successive FPLC purification on HisTrap HP (5 mL, GE Healthcare, 17-1154-01), HiTrap Q HP (5 mL, GE Healthcare, 14-5248-01), and Superdex 200 10/300 GL (GE Healthcare, 17-5175-01) columns. His-tagged protein purification was performed with eluents A (50 mM Tris pH 8, 500 mM NaCl, 20 mM imidazole, 0.5 mM PMSF, and 5% glycerol) and B (as A, except for 200 mM imidazole). After loading and washing with 50 mL of 100% A, a step gradient was applied (50 mL of 20% B, followed by 50 mL of 100% B at a flow rate of 5 mL/min), where our recombinant protein eluted in the second step. Overnight buffer exchange to eluent C (50 mM HEPES pH 8, 25 mM NaCl, and 5% glycerol) was performed using a 15 mL, 20 kDa MWCO Slide-A-Lyzer G3 Dialysis Cassette (Thermo Scientific, A52977). HiTrap Q chromatography was performed using eluents C and D (as C, except for 500 mM NaCl). After loading the dialyzed protein, the column was washed with 25 mL of eluent C and a linear gradient (0-100% D) was applied with a flow rate of 5 mL/min over 10 min. Fractions containing MCM protein were pooled and concentrated 14-fold using a 50 kDa MWCO Amicon Ultra 15 centrifugal filter (Merck, UFC905024) prior to size exclusion chromatography with a buffer containing 50 mM HEPES pH 7.8, 150 mM KCl, 2 mM MgCl_2_, 2 mM TCEP, and 5% glycerol. Samples were concentrated 5-fold with the same centrifugal filters, aliquoted, stored at −80 °C. Protein purity was estimated to be >95% by SDS-PAGE analysis with Coomassie staining (Supplementary Fig. 3) and protein concentration was measured using a Bradford assay.

To measure the effect of malyl-CoA or itaconyl-CoA on MCM activity, we adapted assay conditions described previously^12^, where 0.12 μM MCM (homodimer) were pre-incubated for 10 min with 0.24 μM adenosylcobalamin and 0, 5, 10, 20, or 50 μM malyl-CoA or 0, 1, 3, 6, 14, or 34 μM itaconyl-CoA (concentrations corrected based on purity) in 50 mM HEPES (pH 7.5), 150 mM NaCl, 2 mM MgCl_2_, 2 mM TCEP, and 5% glycerol at 30 °C without agitation in 1.5 mL amber Eppendorf tubes. Methylmalonyl-CoA was added at a final concentration of 65 μM and reactions were quenched 10 min later with 10% formic acid, followed by centrifugation at 16100 *g* for 2 min and filtration of supernatants on 0.2 μm RC filters prior to succinyl-CoA measurement by LC-HRMS/MS using our acyl-CoA method described below.

To determine the effect of malyl-CoA on the absorbance properties of the MCM enzyme, spectra were recorded at 30 °C between 300-800 nm with an AnalytikJena Specord 210 Plus spectrophotometer. A first spectrum was recorded after a 10-min pre-incubation at 30 °C of a reaction mixture (120 μL) containing 12.5 μM MCM (homodimer), 5 μM adenosylcobalamin, 50 mM HEPES (pH 7.5), 150 mM KCl, 2 mM MgCl_2_, 2 mM TCEP, and 5% glycerol^2^. Methylmalonyl-CoA, malyl-CoA, or itaconyl-CoA was then added at a concentration of 500 μM.

### Differential gene expression analysis of 3T3-L1 adipogenesis

Differential gene expression analysis was conducted in R using the deposited mouse 3T3-L1 preadipocytes (Day 0) and differentiated adipocytes (Day 7) dataset [NCBI GEO: GSE20752]^26^. First, quality control analysis was performed using arrayQualityMetrics version 3.46.0 and all data passed quality metrics. Preprocessing steps were taken to background correct, quantile normalize, log-transform, and probe measurements were summarized into single value per gene prior to robust multi-array analysis (RMA) using the affy package version 1.76.0. Differentially expressed genes (DEGs) were identified using the limma package version 3.46.0. P-values were corrected for multiple comparisons using the Benjamini and Hochberg method. The Volcano plot was generated using GraphPad Prism 9.5.0.

### (Pre-)Adipocyte cultivation and differentiation

Murine 3T3-L1 preadipocytes were cultured in DMEM containing 1 g/L glucose, 1 mM pyruvate (Lonza, 12-707F; Capricorn, DMEM-LPXA), 2 mM glutamine (Lonza, BE17-605E; Capricorn, CP22-5004), 10% bovine calf serum (GE Healthcare, HYCLSH30073.03), and 1% penicillin/streptomycin (Life Technologies, 15140122) under 5% CO_2_ at 37 °C prior to differentiation into mature adipocytes^27^. Upon reaching confluency (Day 0), the medium was changed to DMEM containing 1 g/L glucose, 1 mM pyruvate, 2 mM glutamine, 10% fetal bovine serum (FBS; Life Technologies, 10500-064), 1% penicillin/streptomycin, and 1 μg/mL insulin (Sigma-Aldrich, I9278) with an induction cocktail of 1 μM dexamethasone (Sigma-Aldrich, D4902), 500 μM methylisobutylxanthine (Sigma-Aldrich, I5879), and 5 μM troglitazone (Santa Cruz Biotechnology, sc-200904B). On Day 2, this medium was replaced by a medium without the induction cocktail and media refreshed every 2-days thereafter. When used, supplementation with vitamin B12 (Sigma-Aldrich, V6629) began on Day 0 and the concentration is indicated in the figure captions and text, while supplementation with 2 mM itaconate (Sigma-Aldrich, I29204) began on Day 4. Supplementation with 2 mM L-(−)-malate (Sigma-Aldrich, M1000), ^13^C_4_-L-malate (Sigma-Aldrich, 750484), pyruvate (Sigma-Aldrich, P8574), or 1-^13^C-pyruvate (Sigma-Aldrich, 677175) and 2.5 μM vitamin B12 were conducted for 48 h prior to metabolite extraction (Day 12).

### Oil Red O staining

Lipid accumulation was verified using Lonza’s Oil Red O staining for *in vitro* adipogenesis. The Oil Red O stain was prepared by mixing 3 parts of 3 mg/mL Oil Red O (Sigma-Aldrich, O0625) in 99% isopropanol with 2 parts deionized water. Briefly, cells were cultivated in 6-well plates and differentiated, as described above, for 12 days prior to staining. Media was aspirated, cells gently washed with 2 mL PBS, and 2 mL of 10% neutral buffered formalin (G-Biosciences, 786-1056) was added to cells for 60 min. After aspirating the formalin and washing the cells with 2 mL sterile water, 2 mL of 60% isopropanol was added to the cells for 5 min, aspirated, and 2 mL of the Oil Red O stain was added for 5 min. After removing the staining solution, the cells were gently rinsed and stored in tap water until imaging with a compound microscope.

### *Ucp1* gene expression

Pre-adipocytes were cultivated until confluent (Day 0) or until mature (Day 12) in 6-well plates, trypsinized, washed with PBS, and flash-frozen in liquid nitrogen. Samples were stored at −80 °C until RNA isolation using 1 mL Trizol (Thermo Fisher Scientific, 15596026) and 200 μL chloroform, followed by a 5 min incubation on ice. Samples were centrifuged at 13400 x *g* for 15 min at 4°C, and the supernatant was transferred into a new tube. To precipitate the RNA, an equal volume of cold isopropanol and 1 μL glycogen (Merck, 10901393/001) was added to the supernatants and incubated at −20 °C overnight. The samples were vortexed, centrifuged at 13400 x *g* for 10 minutes at 4°C, and the RNA pellets were washed with 800 μL 95% cold ethanol and centrifuged at 14700 x *g* for 10 minutes at 4°C. The washed RNA pellets were air-dried and resuspended in 20 μL of RNase-free water.

RNA aliquots (2 μg) were treated with DNase (Merck, AMPD1) according to the manufacturer’s instructions. To synthesize cDNA, 1 μL of 50 μM oligo(dT) (INVITROGEN, 18418020) and 1 μL of 10 mM dNTP (Promega, U1515) solutions were added to the mixture, incubated at 65 °C for 5 min, and stored on ice prior to the addition of 4 μL of 5x first-strand buffer, 1 μL of 0.1 M DTT, 1 μL of 200 U/μL Superscript III reverse transcriptase (INVITROGEN, 18080044), and 1 μL RNase OUT Recombinant RNase Inhibitor (INVITROGEN, 10777019). Samples were placed in a thermocycler for incubation at 50 °C for 60 min and 70 °C for 15 min.

For real-time qPCR, a reaction mixture (10 μL total volume) was prepared by mixing 5 μL of iQ SYBR Green mix (Bio-Rad, 1708882), 0.3 μL of 10 μM forward primer, 0.3 μL of 10 μM reverse primer, 2 μL of cDNA, and 2.4 μL of RNase-free water. Primer sequences were *Ucp1* forward (ATGACGTCCCCTGCCATTTA), *Ucp1* reverse (GGTGTACATGGACATCGCAC), *Actb* forward (ACTGGGACGACATGGAGAAG), and *Actb* reverse (GTCTCCGGAGTCCATCACAA). Samples were placed in a LightCycler® 480 instrument (Roche) programmed as follows: denaturation at 95 °C for 5 min, 45 cycles at 95 °C for 30 sec, 60 °C for 30 sec, and 72 °C for 15 sec. Relative fold changes in gene expression were calculated using the 2^−ΔΔCt^ Ct method^50^, using beta-actin (*Actb*) as the housekeeping gene.

### LentiCRISPR/Cas9 gene editing

LentiCRISPR V2 plasmids containing the Cas9 coding sequence and our designed control, non-targeting sgRNA (TGCTTTACCGCGTTGGGTAA) or *CLYBL* targeting sgRNA (exon 2: CATAGAGCACTGCTCTCCGG; exon 3: AGACTTTGACCTGGGCACAA; exon 4: CCATTGCAGTCTCCACAAAG) were purchased from Genscript. Plasmids were transformed into OneShot™ *E. coli* Stbl3™ chemically competent cells (Invitrogen, C737303) and purified using a Qiagen midi-prep kit. Viral particles were produced using HEK293 Lenti-X cells, where on Day 0, 5.5 × 10^5^ cells were seeded in 6-well plates containing antibiotic-free growth medium (2.5 mL/well) consisting of DMEM (Gibco, 11995065) containing 4.5 g/L glucose, 1 mM pyruvate, 4 mM glutamate, and 10% FBS. On Day 1, the lentiviral plasmids (500 ng LentiCRISPR V2, 250 ng pMDLg/RRE, 40 ng pMD2.G, and 250 ng pRSV-Rev per well), the transfection reagent TransIT-LT1 (3 μL per well; Sopachem, MIR2300), and OPTI-MEM (12 μL per well; Gibco, 31985062), were combined and incubated for 30 min at room temperature, before adding the mixture dropwise to the packaging cells followed by gentle mixing. After 6 h, the transfection medium was replaced with high-serum medium consisting of DMEM with 30% FBS and 1% penicillin/streptomycin. The spent media of the packaging cells were collected after 72 h and centrifuged at 1500 rpm for 10 min at 4 °C. Following the manufacturer’s protocol, Lenti-X Concentrator (Takara, 63123) was used to concentrate the viral particles 40-fold, prior to resuspension in 600 μL of preadipocyte medium. Eight hours after seeding 6-well plates with 200,000 preadipocytes per well, they were transduced for 48 h with 125 μL of fresh lentivirus and 8 μg/mL polybrene (Sigma-Aldrich, 107689). Polyclonal *CLYBL* knockout cells were generated by co-transfection with two viruses targeting different exons (exon 2 and exon 3 in the KO1 cells; exon 2 and exon 4 in the KO2 cells), while the control line was transfected with the virus containing the non-targeting sgRNA sequence. Cells were selected with 1 μg/mL puromycin (InvivoGen, ant-pr-1) for 5 days, followed by an increase in selection pressure to 3 μg/mL for 4 more days. Frozen stocks were prepared and used for the following experiments. The HsCLYBL-FLAG (Addgene plasmid #111290) and HsCLYBL(D320A)-FLAG (Addgene plasmid #111291) pLys1 expression vectors together with lentiviral packaging plasmids (pMDLg/RRE, pMD2.G, and pRSV-Rev) were used to produce viral particles for generating the *CLYBL* KO+hCLYBL and *CLYBL* KO+hCLYBL^D320A^ rescue cell lines using the same transduction procedure as described above.

### Mitochondrial enrichment and Western blotting analyses

(Pre-)Adipocytes from confluent T175 flasks were trypsinized, washed with PBS, pelleted, and stored at −80 °C. Cell pellets were thawed on ice and mitochondria were isolated by fractionation following a previously described method using a sucrose buffer^51^. Mitochondrial proteins were extracted by freeze-thawing in 50 mM HEPES (pH 7.4) with 50 mM NaCl, 0.5 mM PMSF, 1 mM DTT, and cOmplete Protease Inhibitor Cocktail, separated by electrophoresis on 10% Mini-PROTEAN TGX Precast Protein Gels (Bio-Rad, 4561033, 4561036), and transferred to a PVDF Membrane (Invitrogen, IB24002) using an iBlot2 device. Membranes were blocked with 5% skim milk in TBST for 2 h with agitation at room temperature. Following washing with 5 × 10 mL TBST, primary antibodies were applied overnight with agitation at 4 °C. Membranes were washed with 5 × 10 mL TBST, the fluorescent secondary antibody was applied for 1 h with agitation at room temperature, and the membranes washed with an additional 5 × 10 mL TBST. The primary antibodies used were polyclonal mouse anti-CLYBL (1:1000 dilution; Abnova, Ref: H00171425-B01P, Lot: D528A), polyclonal mouse anti-methylmalonyl-CoA mutase (1:1000 dilution; Abcam, Ref: ab67869, Lot: 1004911-1), and polyclonal rabbit anti-citrate synthetase (1:500 dilution; Abcam, Ref: ab96600, Lot: GR32811462-3). IRDye 680RD Goat Anti-mouse (Li-Cor, Ref: 926-68070, Lot: D10512-15) and IRDye 800CW Goat Anti-rabbit (Li-Cor, 925-32211, Lot: D30110-05) were used as secondary antibodies at a dilution of 1:5000. Western blots were analyzed on a Li-Cor Odyssey FC Imager and ImageJ 1.53K was used for measuring band intensities. Protein quantification was conducted using the Bradford assay^52^.

### Acyl-CoA extraction and measurement by untargeted LC-HRMS/MS

Confluent preadipocytes were differentiated in 10 cm^2^ Petri dishes and on Day 12, acyl-CoA esters were extracted using an adapted solid phase extraction method^28^. After removal of the medium, cells were washed with 10 mL of 0.9% NaCl (37 °C), 750 μL of 3:1 acetonitrile:isopropanol (−20 °C) was added to the cells, and the dish was placed on a metal cold plate (−20 °C). A 250 μL aliquot of 0.1 M KH_2_PO_4_ (pH 6.7, 4°C) containing 0.5 μM ^13^C_3_-malonyl-CoA (Sigma-Aldrich, 655759) as the surrogate standard was added to the plate, cells were scraped, and the cell lysate was transferred to a 2 mL Eppendorf tube. A 700 μL aliquot of 9:3:4 acetonitrile:isopropanol:milli-Q water was added to the dish, the dish was scraped again, and the remaining extract was transferred to the same 2 mL Eppendorf tube. Cell extracts were shaken at 1400 rpm for 10 min (4 °C), centrifuged at 16,100 *g* for 10 min (4 °C), and 1445 μL of the supernatant were transferred to a fresh tube. The metabolite extract was acidified with 360 μL of glacial acetic acid and vortexed prior to acyl-CoA isolation using weak anion exchange solid phase extraction (Supelco, 54127-U). After equilibration of the columns with 1 mL of 9:3:3:4 acetonitrile:isopropanol:milli-Q water:glacial acetic acid and loading of the samples, columns were washed with 1 mL of the equilibration solution, and acyl-CoA esters were eluted with 1.5 mL of 4:1 methanol:250 mM ammonium formate. Metabolite extracts and protein pellets were dried by speedvac overnight at −4 °C.

To improve the quantification of low abundance CoA esters, dried extracts derived from two 10 cm^2^ Petri dishes were pooled during the reconstitution step with 50 μL of 10 mM ammonium carbonate (pH 6.5), before the analysis. Samples were filtered on a 0.2 μm reverse cellulose (RC) filter before injecting 20 μL of the filtrate on a Thermo Vanquish Flex Quaternary LC coupled to a QExactive HF Orbitrap Mass Spectrometer. Acyl-CoA esters were separated on a BEH C18 column (150 × 2.1 mm, 1.7 μm; Waters, 186002353) equipped with a BEH C18 pre-column (2.1 × 5 mm, 1.7 μm; Waters, 186003975) using eluents (A) 10 mM ammonium carbonate (pH 6.5) in Milli-Q water and (B) LC-MS grade acetonitrile; (C) Milli-Q water and (D) 0.2% phosphoric acid were used for column washing^53^. The column was maintained at 60 °C and chromatography was performed at a flow rate of 500 μL/min with the following gradient: 0-2 min, 99% A/1% B; 10 min, 2% A/98% B; 13 min, 2% A/98% B. The flow was diverted to waste and the column was washed at a flow rate of 600 μL/min with the following gradient: 13.1 min, 50% B/50% D; 20 min, 50% B/50% D; 20.1 min, 50% B/50% C; 27 min, 50% B/50% C; 27.1 min, 99% A/1% B; and 31 min, 99% A/1% B. The flow was reduced back to 500 μL/min and re-equilibration of the column was completed with 1 more minute isocratic flow at 99% A/1% B. Electrospray ionization (ESI) was performed in positive mode using a spray voltage of 3.5 kV, capillary temperature of 368 °C, S-lens of 50 V, 56 L/min sheath gas flow, 18 L/min auxiliary gas flow, 2 L/min sweep gas flow, and 460 °C auxiliary gas temperature^53^. Full scan acquisition was performed between 685-1500 *m/z* with a resolution of 60,000, automated gain control (AGC) target of 5e5, and a maximum injection time of 35 ms. Data-dependent acquisition (DDA) was conducted using a resolution of 30,000, AGC target of 1e6, maximum injection time of 100 ms, loop count of 10, isolation window of 0.4 *m/z*, dynamic exclusion of 6 s, and a normalized collision energy of 30 eV. Peak areas were integrated using TraceFinder software (Version 5.1, Supplementary Table 1). Samples were quantified using external calibration curves and the response ratio with the surrogate standard (^13^C_3_-malonyl-CoA) and normalized with mg of total protein (quantified using the Bradford assay). MS^2^ spectral comparison plots were generated in R4.2.1 using the OrgMassSpecR v0.5-3 package. For the stable isotope tracing experiments, FluxFix was used for isotopologue normalization by comparing labeled data with its unlabeled counterpart^54^.

### Untargeted LC-HRMS/MS metabolomics measurements

Preadipocytes were cultured to confluency in 12-well plates and differentiated until Day 12. After washing cells twice with 2 mL of 0.9% NaCl (37 °C), cell metabolism was quenched adding 250 μL of cold (4 °C) methanol:milli-Q water (4:1) containing surrogate standards to the wells and immediately placing the plate on a metal cold plate (−20 °C). After addition of another 30 μL of cold milli-Q water to each well and cell scraping, extracts were transferred to 1.5 mL Eppendorf tubes containing 100 μL of chloroform, and samples were shaken at 1400 rpm for 10 min (4 °C). After addition of 100 μL chloroform and 100 μL milli-Q water (4 °C) to each sample, they were vortexed and centrifuged at 16,100 *g* for 10 min (4 °C) for phase separation. The polar upper phases (100 μL) were transferred to fresh 1.5 mL Eppendorf tubes and metabolite extracts as well as protein interfaces were dried by speedvac overnight at −4 °C. The surrogate standards used were 4-chloro-DL-phenylalanine, 6-chloropurine riboside, 2-chloroquinoline-3-carboxylate, and Nε-trifluoroacetyl-L-lysine. They were added at a concentration of 10 μg/mL to the quenching solution.

Dried extracts were reconstituted in 100 μL of 4:1 acetonitrile:water, filtered on a 0.2 μm reverse cellulose (RC) filter prior to injecting 5 μL of the filtrate on a Thermo Vanquish Flex Quaternary LC coupled to a Thermo Q Exactive HF mass spectrometer. Chromatography was carried out on a SeQuant ZIC-pHILIC 5µm polymer column (150 × 2.1 mm, 5 μm; Supelco, 1.50460.0001) connected to the corresponding SeQuant ZIC-pHILIC Guard pre-column (20 × 2.1 mm; Supelco, 1.50437.0001). The column temperature was maintained at 45 °C. The flow rate was set to 0.2 mL/min and the mobile phases consisted of (A) 20 μM ammonium carbonate in water, pH 9.2, containing 0.1% InfinityLab deactivator additive (Agilent) and (B) acetonitrile. The following gradient was applied: 0 min, 80% B; 3 min, 80% B; 18 min, 20% B; 19 min 20% B; 20 min 80% B; 24.5 min 80% B; 25.5 min 80% B (0.4 mL/min); 29.5 min 80% B (0.4 mL/min); 30 min 80% B. All MS experiments were performed using electrospray ionization with polarity switching enabled (+ESI/-ESI). The following source parameters were applied: sheath gas flow rate, 25; aux gas flow rate, 15; sweep gas flow rate, 0; spray voltage, 4.5 kV (+) / 3.5 kV (–); capillary temperature, 325 °C; S-lens RF level, 50; aux gas heater temperature, 50 °C. The Orbitrap mass analyzer was operated at a resolving power of 30,000 in full-scan mode (scan range: *m/z* 75-1000; automatic gain control target:1e6; maximum injection time: 240 ms). MS^2^ spectra were acquired with DDA using the same spray settings but without polarity switching, and the following parameters: resolution of 30,000, AGC target of 1e5, maximum injection time of 50 ms, loop count of 5, isolation window of 4 *m/z*, dynamic exclusion of 10 s, and a normalized collision energy of 30 eV. Data were acquired with Thermo Xcalibur software (Version 4.3.73.11) and analyzed with TraceFinder (Version 5.1, Supplementary Table 2). The dataset was normalized to the signal intensity (peak area) of surrogate standard Nε-trifluoroacetyl-L-lysine.

### Extracellular propionate measurements

Two-day old spent medium was collected on Day 12 of adipogenesis, RC filtered, and stored at −80 °C until analysis. Samples were thawed and 180 μL aliquots of spent media were transferred to 1.5 mL Eppendorf tubes containing 20 μL of 2-ethylbutyric acid (surrogate standard) and 10 μL of 37% HCl, followed by shaking at 1400 rpm for 15 min at 15°C. A 1 mL aliquot of ethyl ether was added to the samples, followed by identical shaking. Samples were centrifuged at 21,000 *g* for 5 min at 15 °C and 900 μL extract (upper phase) was transferred to a fresh 2 mL Eppendorf tube. After addition of another 1 mL aliquot of ethyl ether to the remaining lower phase, samples were shaken for 5 min, centrifuged, and 900 μL supernatant was pooled with the first extract. A 250 μL aliquot of final extract was transferred to a GC-MS vial containing 25 μL of MTBSTFA with 1% tert-Butyldimethylchlorosilane (TBDMSCI). Vials were capped using a crimper, vortexed, and samples were left to derivatize for at least 2 h before injection. GC-MS analysis was performed with an Agilent 8890 GC – 5977B MS instrument. A sample volume of 1 μL was injected into a Split/Splitless inlet, operating in split mode (20:1) at 280 °C. The gas chromatograph was equipped with a ZB-5Msplus capillary column (30□ m, I.D. 250 μm, film 0.25 μm; Phenomenex, 7HG-G030-11-GGA) and a 5 m GUARDIAN column in front of the analytical column. Helium was used as carrier gas with a constant flow rate of 1.4□ mL/min. The GC oven temperature was held at 80 °C for 1 min and increased to 170 °C at 10 °C/min, followed by a post-run time at 280 °C for 5 min. The total run time was 15 min. The transfer line temperature was set to 280 °C. The mass selective (MS) detector was operating under electron ionization at 70 eV. The MS source was held at 230 °C and the quadrupole at 150 °C. The detector was switched off during the elution of MTBSTFA. For precise quantification, GC-MS measurements of the compounds of interest were performed in selected ion monitoring mode (Supplementary Table 3). Acquired GC-MS data were processed using MassHunter Workstation Software Quantitative Analysis (Version 10.2 / Build 10.2.733.8). Target compounds were identified by retention time and ion ratios using an in-house mass spectral library. The data set was normalized by using the response ratio of the integrated peak area of the target compound and the integrated peak area of the internal standard. Absolute concentrations were determined using an external calibration curve made of an authentic standard mixture (Merck, CRM46975).

### Targeted LC-MS/MS adenosylcobalamin measurements

Confluent adipocytes cultured in 6-well plates were extracted on differentiation Day 12 to measure adenosylcobalamin levels. After removal of spent media by aspiration, cells were washed with 10 mL of 0.9% NaCl (37 °C) and quenched by adding 600 μL of methanol (−20 °C) and placing the dish immediately on a cold plate (−20 °C). After adding another 600 μL aliquot of 50% methanol (4 °C), cells were scraped and resulting extracts were split in 2 × 1.5 mL amber Eppendorf vials that were prefilled with 450 μL of chloroform (4 °C). Samples were shaken at 1400 rpm for 10 min at 4 °C and centrifuged at 16100 *g* for 10 min at 4 °C for phase separation. A 750 μL aliquot of the aqueous phase from each vial was transferred to a fresh amber Eppendorf tube, and samples were dried overnight by speedvac (−4 °C) prior to storage at −80 °C until analysis. The non-polar chloroform phases were discarded, and protein interfaces derived from each culture dish were combined, dried overnight, and used for protein normalization.

Dried polar metabolite extracts were resuspended in 100 μL of 20 mM ammonium formate, pH 3.5, Acetonitrile (4:1) containing ^13^C_3_-caffeine (0.1 μg/mL) as internal standard. LC-MS analyses were performed on an Exion LC coupled to a 7500 Triple Quad MS instrument (SCIEX) equipped with an Optiflow Pro Ion Source,operated in electrospray ionization mode. Chromatography was performed on an ACQUITY UPLC CSH C18 column (2.1 × 100 mm, 1.7 μm; Waters, 186005297) protected by a VanGuard pre-column (2.1 × 5 mm; Waters, 186003975) maintained at 40 °C. The autosampler was kept at 4 °C and a sample injection volume of 2 μL was used. The flow rate was set to 0.2 mL/min for the entire 18-min run. Mobile phases consisted of (A) 20 mM ammonium formate, pH 3.5, and (B) 100% Acetonitrile. After 2 min isocratic delivery at 2% B, a linear gradient to 98% B over 7 min was used to elute all target components. Following 3 min isocratic delivery at 98% B, a re-equilibration phase on starting conditions (2% B) was applied for 6 min. Target compounds were measured in multiple reaction monitoring mode. The source and gas parameters applied were as follows: ion source gas 1 and 2 were maintained at 35 psi and 70 psi, respectively; curtain gas was at 40 psi; CAD gas was at 8 psi; source temperature was held at 350 °C. In positive ESI mode, the spray voltage was set to 2000 V and in negative ESI mode, it was set to 1500 V. Mass spectrometric data were acquired with SCIEX OS (Version 3.0.0) and analyzed with MultiQuant (Version 3.0.3). Target compounds were identified by retention time and ion ratio (Supplementary Table 4). The dataset was normalized by using the response ratio of the integrated peak area of target compound and the integrated peak area of the internal standard (^13^C_3_-caffeine) and mg of total protein (quantified using the Bradford assay).

### Statistical analyses

Kinetic parameters for all enzymes were estimated in GraphPad Prism (9.5.0) using the non-linear Michaelis-Menten fit model. Assuming normal distribution, an ordinary one-way analysis of variance (ANOVA) using Fisher’s LSD *posthoc* tests were performed in GraphPad Prism 10.2.3, which was also used to construct histograms and line plots. The data was autoscaled prior to heatmap generation using MetaboAnalyst 6.0.

## Acknowledgements

This work was supported by the Luxembourg National Research Fund (FNR) through an AFR Postdoc grant (No. 9180195) to K.W.E., the University of Luxembourg through Internal Research Project (IRP; project *HYMEPI*) funding to C.L.L., and the European Union’s Horizon 2020 Research and Innovation Programme through grant No. 814418 (*SinFonia*) to C.L.L. We thank Emile Van Schaftingen, Guido Bommer, and Toshimori Kitami for fruitful discussions related to this work, in addition to Lasse Sinkkonen for kindly providing the murine 3T3-L1 preadipocytes. We thank Dominic Esposito for providing the pDEST527 vector (Addgene plasmid #1151) and Vamsi Mootha for providing the *Pseudomonas aeruginosa* succinyl-CoA:itaconate CoA transferase (PaIct) (Addgene plasmid #111293), HsCLYBL-FLAG (Addgene plasmid #111290), and HsCLYBL(D320A)-FLAG (Addgene plasmid #111291) plasmids. Finally, we thank the Luxembourg Centre for Systems Biomedicine Metabolomics and Lipidomics Platform (RRID:SCR_024769) for running metabolomics samples, and Lucia Gallucci and Xiangyi Dong for their technical and analytical support.

## Ethics & Inclusion Statement

Not applicable to this article.

## Data Availability Statement

The raw untargeted metabolomics data associated with this manuscript is available on GNPS (https://doi.org/doi:10.25345/C5G44J284).

## Code Availability Statement

Custom code was not used in this article.

---

**Extended Data Figure 1.**
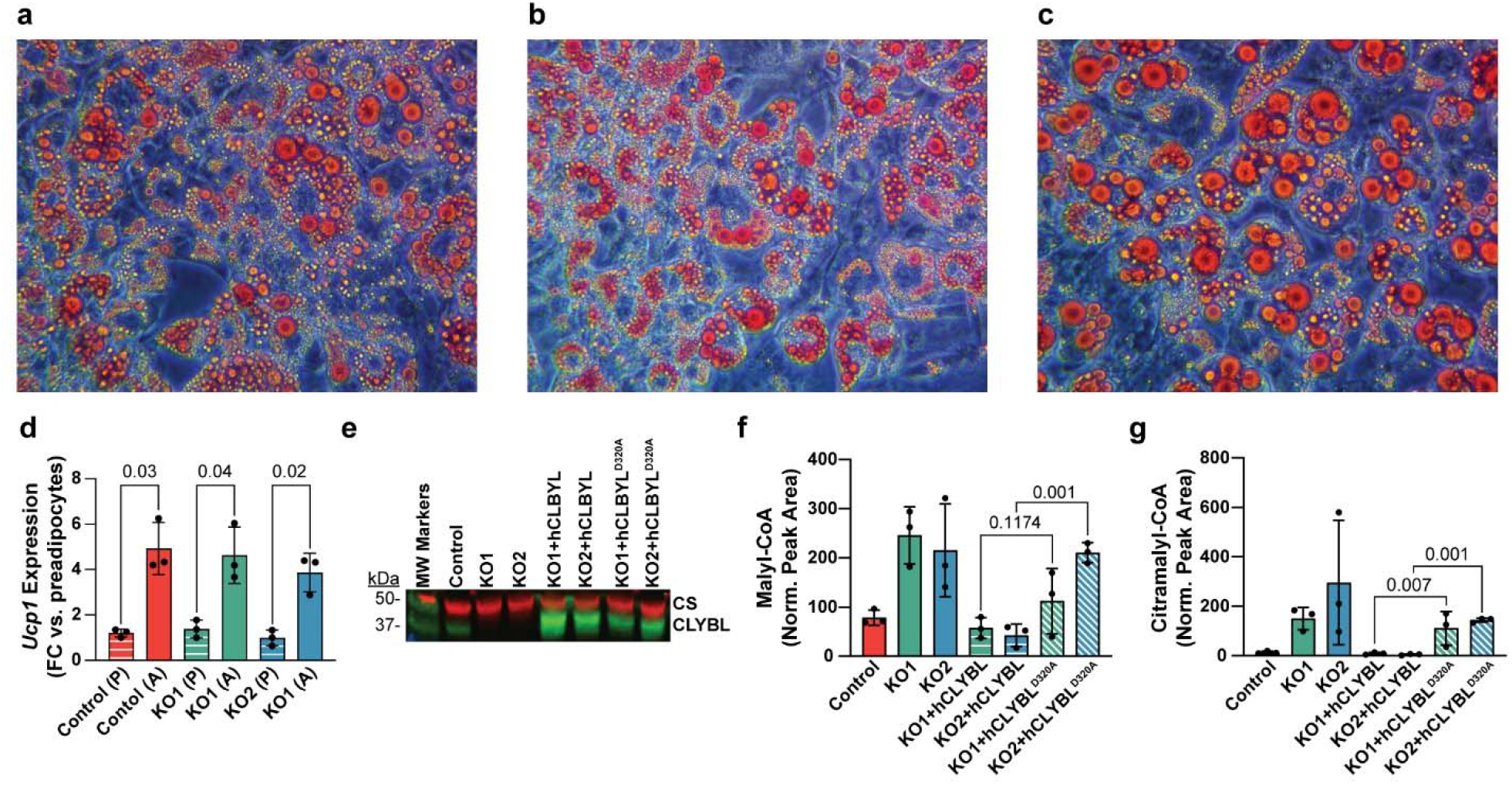
Characterization of 3T3-L1 adipogenesis and rescue experiments in *CLYBL* KO cells. Oil Red O staining and imaging (40x magnification) of mature 3T3-L1 control (**a**), *CLYBL* KO1 (**b**), and *CLYBL* KO2 (**c**) adipocytes (Differentiation Day 12). (**d**) Relative mRNA expression of the brown adipocyte marker uncoupling protein 1 (*Ucp1*) measured by qPCR in control and *CLYBL* KO preadipocytes (P; striped bars) and 12-Day mature adipocytes (A; solid bars). Fold changes in mature adipocytes versus preadipocytes were calculated using the 2^−ΔΔCt^ method, using *Actb* as the reference gene. Data are means ± SD (n = 3) and statistical significance was determined using the Welch t-test. Rescue experiments were performed with *CLYBL* KO1 and KO2 cell lines transduced with FLAG-tagged human CLYBL (hCLYBL) or a catalytically defective variant thereof (hCLYBL^D320A^). (**e**) Western blot analysis of mitochondrial fractions (38 μg protein/lane) of mature adipocytes for detection of CLYBL (35 kDa), hCLYBL-FLAG (39 kDa), hCLYBL^D320A^-FLAG (39 kDa), and CS (49 kDa, loading control) showed that CLYBL expression was successfully restored. Malyl-CoA (**f**) and citramalyl-CoA (**g**) accumulation in *CLYBL* KO cells was reversed by expressing active, but not inactive CLYBL. Data are means ± SD (n = 3), statistically significant changes in *CLYBL* KO+hCLYBL vs. *CLYBL* KO+hCLYBL^D320A^ cells were determined with an ANOVA followed by Fisher’s LSD *posthoc* test. Norm. Peak Area corresponds to data normalized to total protein). CLYBL, citrate lyase beta-like protein; CS, citrate synthase; FC, fold change.

**Extended Data Figure 2.**
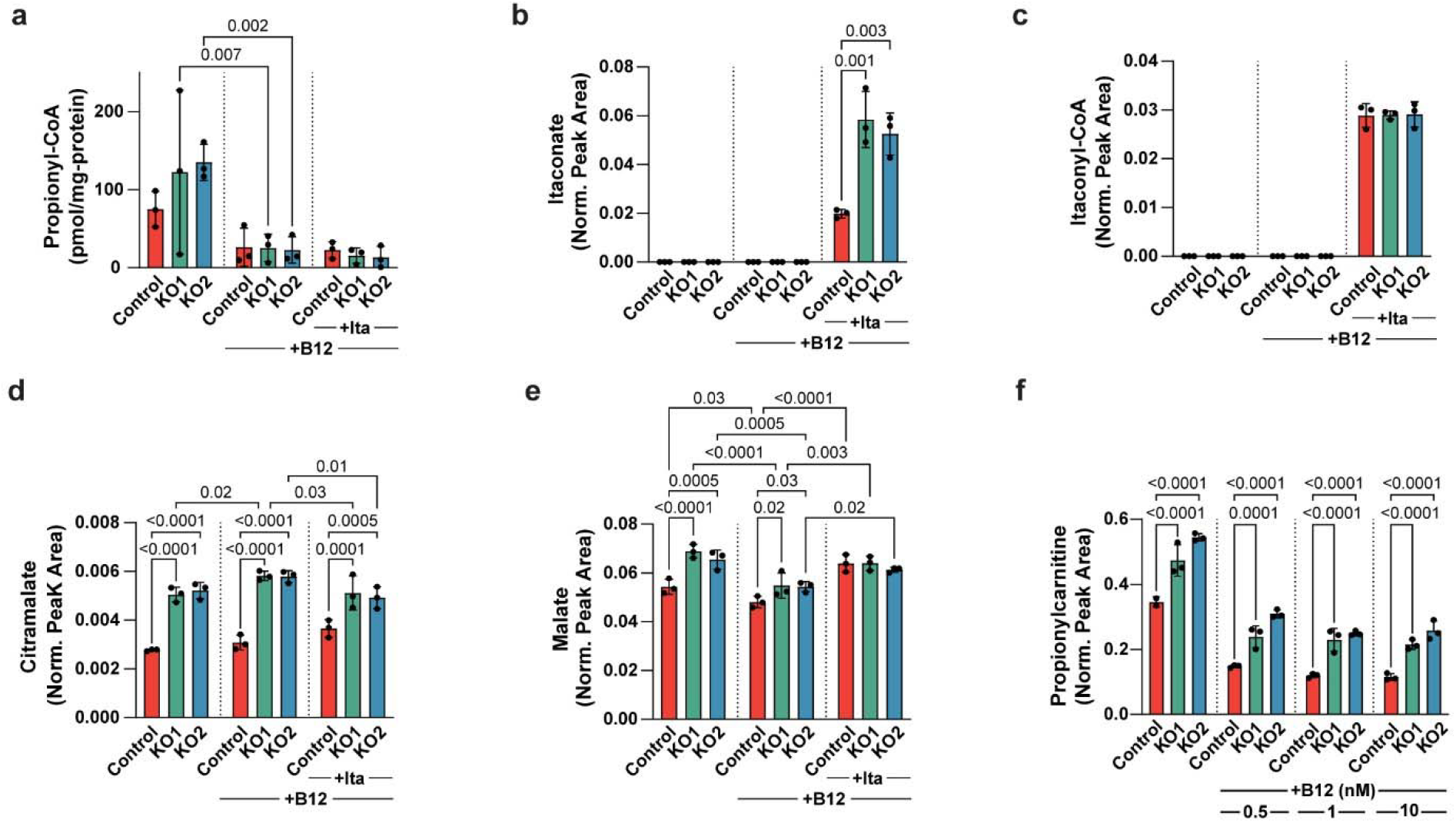
Propionate metabolism perturbation caused by *CLYBL* gene deletion persists upon B12 supplementation but is not exacerbated by itaconate supplementation. Levels of (**a**) propionyl-CoA, (**b**) itaconate, (**c**) itaconyl-CoA, (**d**) citramalate, and (**e**) malate in extracts of control and *CLYBL* KO 3T3-L1 adipocytes on differentiation Day 12, untreated or supplemented with 1 nM B12 (+B12; from Day 0) and/or 2 mM itaconate (+Ita; from Day 4). (**f**) Effect of B12 supplementation at increasing concentration (from Day 0) on propionylcarnitine levels. The values shown are means ± SD (n = 3) and statistical significance was determined through ANOVA followed by Fisher’s LSD *posthoc* test. Acyl-CoA esters were quantified using the ion ratio with the surrogate standard (^13^C_3_-malonyl-CoA) and external calibration curves and normalized by total protein, except itaconyl-CoA where Norm. Peak Area corresponds to the peak area normalized with the surrogate standard peak area and total protein. For other metabolites, Norm. Peak Area corresponds to data normalized to the surrogate standard (Nε-trifluoroacetyl-L-lysine) only.

