## Supplemental Figures 1-6 & Tables 1-4 for "CLYBL averts methylmalonyl-CoA mutase inhibition and loss of vitamin B12 by repairing malyl-CoA"

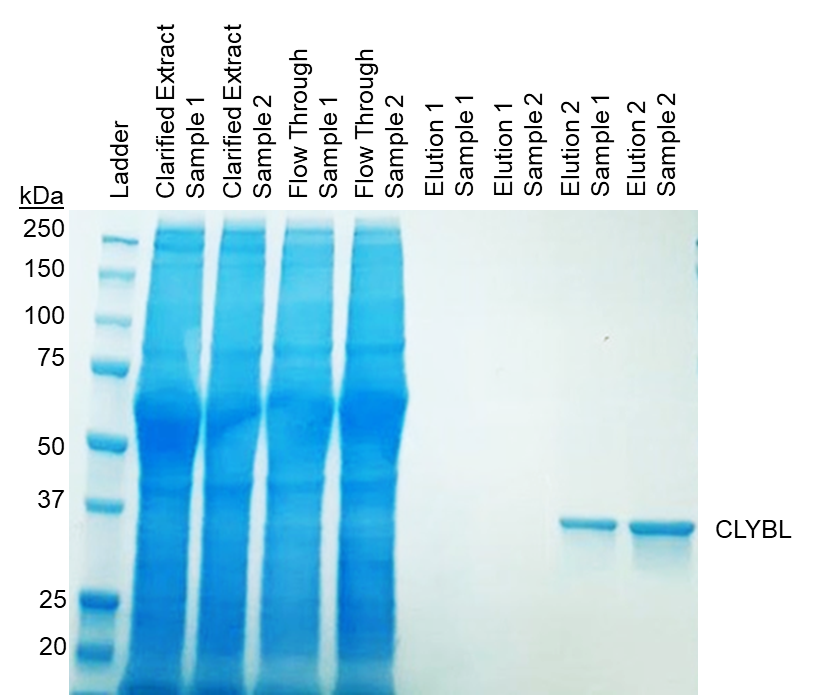

**Supplementary Fig. 1 |** Representative SDS-PAGE analysis with Coomassie staining of affinity purification fractions of recombinant human CLYBL from two lysates (sample 1 and sample 2) of HEK293FT cells overexpressing the FLAG-tagged protein (predicted MW of the full-length CLYBL fusion protein = 40 kDa; GenBank ID AAH34360.1). Elution 1 and Elution 2 correspond to the first and subsequent pooled fractions, respectively, eluted with the FLAG peptide. The protein concentration of the pooled Elution 2 fractions corresponded to 70 μg/mL and 163 μg/mL for sample 1 and sample 2, respectively, and 32 μL of this pool was loaded in the indicated lanes.

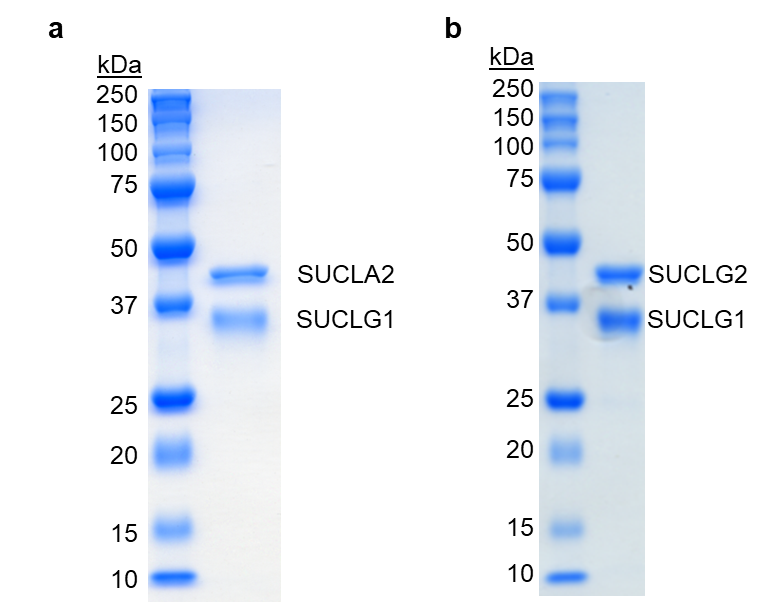

**Supplementary Fig. 2 |** Representative SDS-PAGE analysis with Coomassie staining of the purified recombinant human (**a**) ADP-forming and (**b**) GDP-forming succinyl-CoA ligase (SUCL) preparations used in this study. SUCL was produced in *E. coli* BL21 cells co-transformed with *SUCLG1* (ADP/GDP-forming subunit alpha, predicted MW = 34 kDa; NCBI RefSeq NP_003840.2, corresponding to amino acids 28-333) and *SUCLA2* (ADP-forming subunit beta, predicted MW = 46 kDa; NCBI RefSeq NP_003841.1, corresponding to amino acids 53-463) or *SUCLG2* (GDP-forming subunit beta, predicted MW = 44 kDa; NCBI RefSeq NP_003839.2, corresponding to amino acids 38-420) followed by nickel affinity purification of the polyhistidine-tagged proteins. The protein concentration of the final purified fraction was 1.7 mg/mL for the ADP-forming complex and 1.1 mg/mL for the GDP-forming complex.

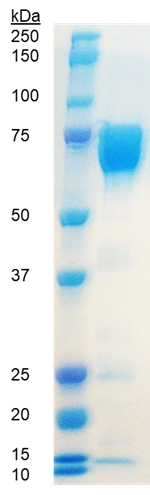

**Supplementary Fig. 3** | Representative SDS-PAGE analysis with Coomassie staining of the purified recombinant human MCM preparation used in this study (predicted MCM fusion protein MW = 80 kDa; NCBI RefSeq NP_000246.2, corresponding to amino acids 34-750). The recombinant His-tagged protein was overexpressed in *E. coli* BL21 cells and purified by nickel affinity chromatography (HisTrap HP) followed by strong anion exchange (HiTrap Q HP) and size exclusion chromatography. The protein concentration of the final purified fraction was 4.7 mg/mL and 15 μL of a 10-fold diluted aliquot was loaded on the gel.

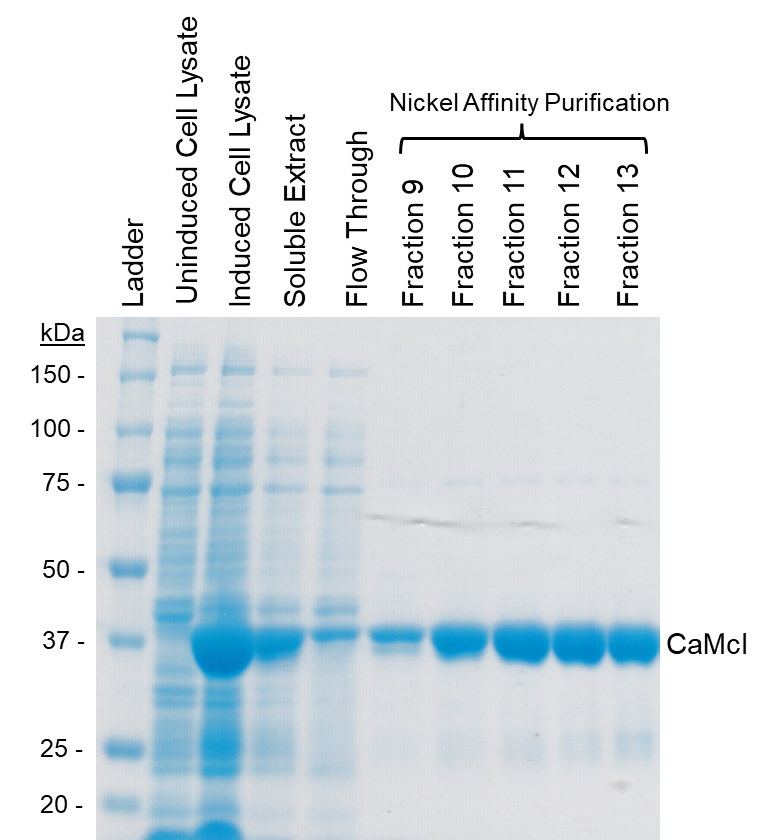

**Supplementary Fig. 4 |** Representative SDS-PAGE analysis with Coomassie staining showing the purification of the recombinant *Chloroflexus aurantiacus* malyl-CoA/citramalyl-CoA lyase (CaMcl) used in this study to produce malyl-CoA and citramalyl-CoA. CaMcl was overexpressed as a His-tagged protein in *E. coli* RosettaBlue(DE3) cells and purified by nickel affinity chromatography (predicted MW = 41 kDa; GenBank ID AGR55786.1). Active fractions were pooled, desalted, and 10% glycerol was added prior to storage at -80 °C.

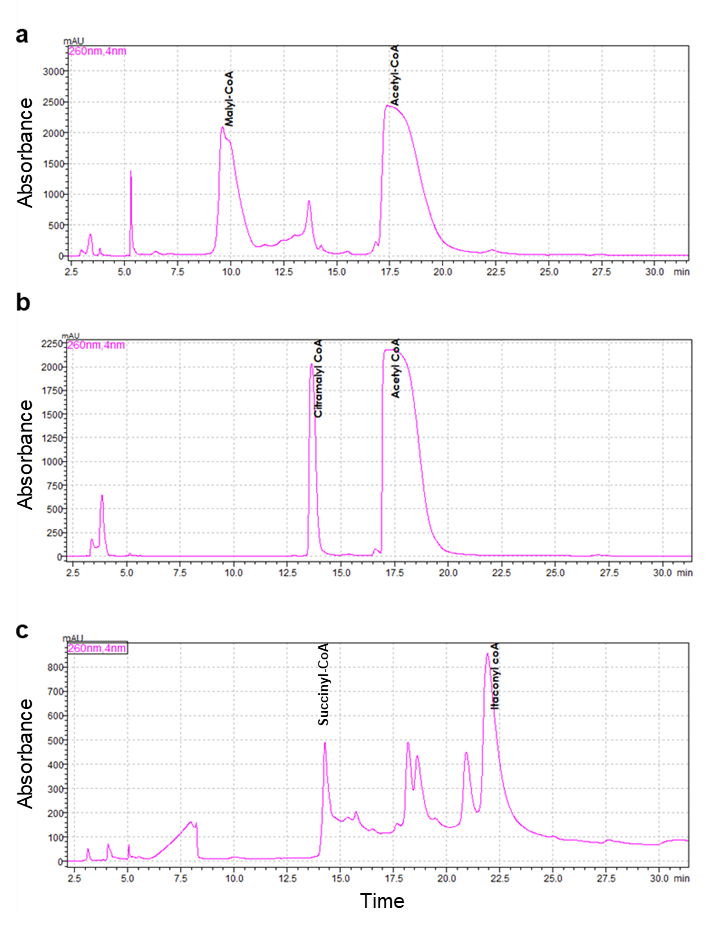

**Supplementary Fig. 5** | Representative chromatograms (λ = 260 nm) of the semi-preparative HPLC purifications of acyl-CoA esters synthesized for this study. *Chloroflexus aurantiacus* malyl-CoA/citramalyl-CoA lyase (CaMcl) was used to produce (**a**) malyl-CoA and (**b**) citramalyl-CoA from acetyl-CoA and glyoxylate or pyruvate, respectively. (**c**) Itaconyl-CoA was synthesized from succinyl-CoA and itaconate, using *Pseudomonas aeruginosa* succinyl-CoA:itaconate CoA transferase (PaIct). Relevant peak fractions were collected and the identity of the purified acyl-CoAs was confirmed using high-resolution mass spectrometry (MS^1^ and MS^2^).

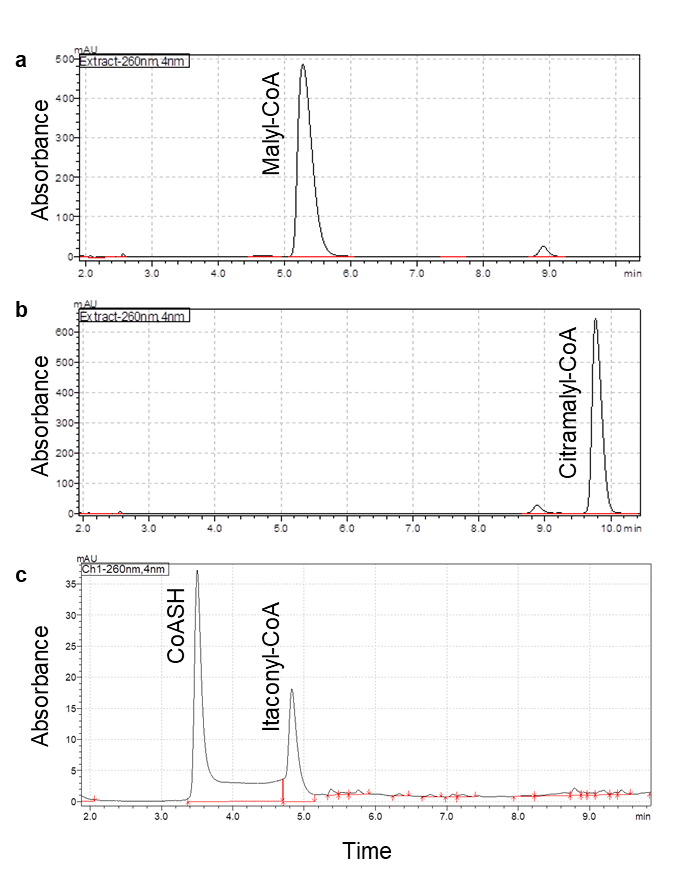

**Supplementary Fig. 6 |** Representative analytical HPLC chromatograms (λ = 260 nm) showing the purity of our synthesized and purified acyl-CoAs: (**a**) malyl-CoA (97% pure, RT = 5.3 min), (**b**) citramalyl-CoA (96% pure, RT = 9.8 min), (**c**) itaconyl-CoA (27% pure; RT = 4.8 min; free CoASH, RT = 3.5 min).

**Supplementary Table 1 |** Retention times, exact masses ([M+H]^+^), and MS^2^ fragments of acyl-CoA esters detected with our BEH C18, untargeted LC-HRMS/MS method. ^13^C_3_-Malonyl-CoA was used as the surrogate standard.

| **Compound** | **Retention Time (Min)** | **[M+H]^+^** | **MS^2^ Fragments** | **Molecular Formula** | **PubChem CID** |
| --- | --- | --- | --- | --- | --- |
| ^13^C_3_-Malonyl-CoA | 1.40 | 857.1336 | 350.1369;305.1439;136.0618;428.0371;159.0587 |  |  |
| Malonyl-CoA | 1.40 | 854.1234 | 347.1271;428.0368;303.1372;245.0590;136.0619 | C24H38N7O19P3S | 644066 |
| Malyl-CoA | 1.42 | 884.134 | 377.1373;428.0364;275.0692;136.0617;330.0592 | C25H40N7O20P3S | 440302 |
| Succinyl-CoA | 1.80 | 868.1391 | 361;1429;428.0370;259.0749;136.0620;330.0600 | C25H40N7O19P3S | 92133 |
| Citramalyl-CoA | 2.03 | 898.1497 | 391.1530;428.0363;289.0849;136.0619;330.0602 | C26H42N7O20P3S | 11966212 |
| Methylmalonyl-CoA | 2.10 | 868.1391 | 361.1422;317.1525;428.0363;259.0744;136.0615 | C25H40N7O19P3S | 123909 |
| Coenzyme A | 2.25 | 768.123 | 261.1266;428.0366;159.0587;136.0618;341.0929 | C21H36N7O16P3S | 87642 |
| Glutaryl-CoA | 2.78 | 882.1547 | 375.1577;136.0616;273.0895;428.0359;159.0588 | C26H42N7O19P3S | 3081383 |
| 3-Hydroxy-3-methylglutaryl-CoA | 3.20 | 912.1653 | 405.1686;428.0363;303.1007;177.0697;330.0589 | C27H44N7O20P3S | 91506 |
| Itaconyl-CoA | 3.60 | 880.1391 | 373.1427;428.0368;271.048;136.0621;261.1274 | C26H40N7O19P3S | 11966125 |
| Acetyl-CoA | 4.90 | 810.1336 | 303.1372;428.0367;201.0692;136.0619;330.0595 | C23H38N7O17P3S | 444493 |
| Acetoacetyl-CoA | 4.92 | 852.1441 | 345.1474;428.0363;243.0795;261.1263;159.0585 | C25H40N7O18P3S | 92153 |
| 3-Hydroxybutyryl-coenzyme A | 4.96 | 854.1598 | 347.1631;428.0364;245.0953;136.0617;330.0593 | C25H42N7O18P3S | 644065 |
| Dephospho-Coenzyme A | 5.06 | 688.1567 | 261.1261;136.0616;159.0584;348.0694;243.1156 | C21H35N7O13P2S | 444485 |
| Propionyl-CoA | 5.21 | 824.1492 | 317.1529;428.0367;215.0849;477.0574;136.0617 | C24H40N7O17P3S | 92753 |
| Crotonoyl-CoA | 5.37 | 836.1492 | 329.1519;227.0843;428.0356;227.0843;136.0616 | C25H40N7O17P3S | 5497143 |
| Butyryl-CoA | 5.52 | 838.1649 | 331.1685;428.0367;229.1006;136.0618;330.0594 | C25H42N7O17P3S | 122283 |
| Benzoyl-CoA | 5.77 | 872.1493 | 365.1525;428.03623;263.0845;136.0617;330.0593 | C28H40N7O17P3S | 9543169 |
| Oleoyl-CoA | 9.54 | 1032.368 | 525.3718;136.0618;428.0363;261.1263;159.0586 | C39H68N7O17P3S | 5497111 |
| Stearoyl-CoA | 10.00 | 1034.384 | 527.3874;136.0618;428.0363;236.1264;159.0596 | C39H70N7O17P3S | 94140 |

**Supplementary Table 2 |** Retention times, exact masses (*m/z*), and MS^2^ fragments of metabolites detected with our zic-pHILIC, untargeted LC-HRMS/MS method. Nε-trifluoroacetyl-L-lysine as used as the surrogate standard.

| **Compound** | **Retention Time (Min)** | ***m/z*** | **Ion** | **MS^2^ Fragments** | **Molecular Formula** | **PubChem CID** |
| --- | --- | --- | --- | --- | --- | --- |
| Nε-trifluoroacetyl-L-lysine | 3.12 | 243.0951 | [M+H]^+^ | 180.0632;197.0898;243.0951 | C8H13F3N2O3 | 2802360 |
| Propionyl-carnitine | 3.37 | 218.1392 | [M]^+^ | 85.0285;216.1388;159.0653;60.0810 | C10H20NO4^+^ | 107739 |
| Methylmalonate | 10.76 | 117.0188 | [M-H]^-^ | 116.9287;73.0293;99.9259;117.9288 | C4H6O4 | 487 |
| Citramalate | 11.8 | 147.0293 | [M-H]^-^ | 87.0088;147.0300;85.0295;129.0195 | C5H8O5 | 1081 |
| Itaconate | 11.88 | 129.0187 | [M-H]^-^ | 85.0294;129.0194;86.0328;52.3314 | C5H6O4 | 811 |
| Succinate | 12.19 | 117.0188 | [M-H]^-^ | 73.0293;117.0194;99.0088;74.0326 | C4H6O4 | 487 |
| Malate | 12.96 | 133.0137 | [M-H]^-^ | 115.0037;71.0136;133.0143;72.9929 | C4H6O5 | 525 |

**Supplementary Table 3 |** Quantification and qualification ions (m/z) and dwell times for the GC-MS-based measurement of extracellular short chain fatty acids derivatized with MTBSTFA. Propionic acid and 2-ethylbutyric acid had retention times of 4.6 and 7.5 min, respectively.

| **Derivatives** | **Quant-Ion** | **Qual-Ion** | **Qual-Ion** | **Dwell Time** |
| --- | --- | --- | --- | --- |
|  | **(*m/z*)** | **(*m/z*)** | **(*m/z*)** | **(ms)** |
| Propionic acid 1TBDMS | **131.1** | 75 | 115 | 20 |
| 2-Ethylbutyric acid 1TBDMS | **173.1** | 75 | 115 | 20 |

**Supplementary Table 4 |** MRM parameters used for the targeted LC-MS/MS measurements of intracellular adenosylcobalamin. Adenosylcobalamin and ^13^C_3_-caffeine had retention times of 5.6 and 5.7 min, respectively.

| **Polarity** | **Compound** | **Q1 mass (Da)** | **Q3 mass (Da)** | **Dwell time (ms)** | **EP (V)** | **CE (V)** | **CXP (V)** | **Q0D (V)** |
| --- | --- | --- | --- | --- | --- | --- | --- | --- |
| POS | AdoCobalamin_(2+)_QUAN_1 | 790.5 | 665.53 | 15 | 10 | 30 | 36 | -10 |
| POS | AdoCobalamin_(2+)_QUAL_1 | 790.5 | 147.109 | 15 | 10 | 129 | 14 | -10 |
| POS | IS1_CAF_QUAN | 198.1 | 140.016 | 15 | 10 | 26 | 13 | -10 |
| POS | IS1_CAF_QUAL | 198.1 | 112.025 | 15 | 10 | 30 | 6 | -10 |

Settling time 15 ms, Pause time 5 ms
